# Autophagy selectively clears ER in inflammation-induced muscle atrophy

**DOI:** 10.1101/2025.05.06.650963

**Authors:** Ursula K. Dueren, Alan An Jung Wei, A. Elisabeth Gressler, Oliver Popp, Paolo Grumati, Matthias Selbach, Anna Katharina Simon, Thomas Sommer

## Abstract

Skeletal muscle atrophy is a pathological condition characterized by the progressive loss of muscle mass and function, driven by factors such as disuse, inflammation, and aging. While the ubiquitin-proteasome system is established as the central mediator of myofibrillar protein degradation, the role of autophagy in selective protein turnover remains largely unexplored. To address this, we employed a quantitative, time-resolved analysis of protein synthesis and degradation in C2C12 myotubes undergoing TNF-α-induced atrophy, using dynamic Stable Isotope Labeling by Amino Acids in Cell Culture (dynamic SILAC) coupled with LC-MS/MS. Our data challenges the classical view of atrophy as a uniform, degradation-centric process. Instead, we reveal temporally distinct patterns of selective protein turnover, including differential degradation of myofibrillar, ribosomal, and endoplasmic reticulum (ER)-resident proteins. Early atrophy is characterized by suppressed short-term protein synthesis, increased ubiquitin-ligase expression, proteasomal activation, and ribosome turnover. In contrast, late atrophy features proteasome-dependent myofibrillar protein degradation, selective synthesis of mitochondrial ribosomes and cytoplasmic ribosome degradation, indicative of metabolic adaptation. Moreover, we identify a temporal shift in autophagic selectivity: from ER homeostasis maintenance to a stress-induced ER-degradation program. Notably, inhibition of autophagy during atrophy leads to accumulation of ER-phagy receptors Tex264 and Calcoco1, implicating ER-phagy as a key contributor to atrophic remodeling, underscoring an underexplored regulatory mechanism in muscle proteostasis. By elucidating the role of autophagy in degradation of the ER, this study opens new avenues for therapeutic interventions targeting proteostasis regulation in inflammation-induced muscle-wasting disorders, ultimately contributing to a more refined understanding of muscle atrophy beyond proteasomal degradation.

## Introduction

Skeletal muscle atrophy is a pathological condition characterized by the loss of muscle mass and strength, driven by diverse stimuli including disuse, malnutrition, inflammation, and aging. This process has profound implications for overall health, leading to impaired mobility, metabolic dysregulation, and increased disease susceptibility. Muscle atrophy has been attributed to an imbalance in proteostasis, where protein degradation surpasses synthesis, primarily affecting myofibrillar proteins and cellular organelles (Sandri, 2013; Bodine and Baehr, 2014). This long-standing paradigm is supported by several observations. First, the majority of muscle proteins are myofibrillar, and they undergo significant breakdown during atrophying conditions alongside an activation of proteolytic systems (Wing and Goldberg, 1993; Mitch *et al*., 1994). Additionally, suppressed protein synthesis via the Akt/mTOR pathway and activation of Forkhead-Box-Protein (FoxO) signaling further contribute to muscle loss by upregulation of E3 ubiquitin ligases such as Muscle RING-finger protein-1 (MuRF1) and Atrogin-1, promoting ubiquitin proteasome system (UPS)-dependent degradation of sarcomeric proteins (Bodine *et al*., 2001; Bodine and Baehr, 2014). Despite that, denervation model studies in rats have shown a more nuanced process with both increased protein degradation and elevated protein synthesis rates (Argadine *et al*., 2009). This complexity underscores the need for a time-resolved approach to observe protein dynamics across different atrophy-inducing conditions, fiber-type compositions and model organisms, rather than assuming a uniform degradation-centric model.

C2C12 mouse skeletal muscle myotubes are widely used as a skeletal muscle model, allowing for a controlled investigation of atrophy-inducing stimuli such as pro-inflammatory cytokines like tumor necrosis factor-alpha (TNF-α). This leads to activation of multiple pathways including FoxO, nuclear factor kappa B (NF-κB) and transcription factor EB (TFEB) signaling, encompassing protein degradation and autophagy (Li and Reid, 2000; Cai *et al*., 2004; Dogra *et al*., 2007; Keller *et al*., 2011; De Larichaudy *et al*., 2012; Bernacchioni *et al*., 2021). TNF-α-driven inflammation is particularly relevant in sarcopenia (Lees, Zwetsloot and Booth, 2009; Sishi and Engelbrecht, 2011; Pascual-Fernández *et al*., 2020) and chronic muscle-wasting diseases, like cachexia (Oliff *et al*., 1967; Beutler, 1985) where it exacerbates proteolytic activity and disrupts proteostasis.

Most studies focus on the influence of the UPS on atrophy progression whereas other proteolytic systems like the autophagy/lysosome pathway – primarily referring to macroautophagy, distinct from chaperone-mediated autophagy (CMA) and microautophagy – has emerged as a critical but still underexplored regulator of muscle proteostasis. Hereafter, for simplicity, we refer to macroautophagy as “autophagy”. Autophagy plays a central role in the selective degradation of damaged proteins and organelles through the formation of double-membrane vesicles (autophagosomes) that engulf and thereby deliver substrates to lysosomes for degradation. Whilst basal autophagy is essential for cellular homeostasis, autophagy is upregulated upon a variety of atrophying stimuli like inflammation-induced atrophy (Keller *et al*., 2011; Bernacchioni *et al*., 2021) and denervation (O’Leary and Hood, 2009). Suppression of autophagy – shown by a muscle-specific deletion of a key autophagy gene, *Atg7* – results in muscle atrophy, accompanied by disorganization of the sarcomere, abnormal mitochondria, sarcoplasmic reticulum distension and formation of aberrant membranous structures (Masiero *et al*., 2009). On the other hand, several studies suggest that activation of autophagy during catabolic conditions can aggravate muscle loss. Loss-of-function mutations in Jumpy – a phosphatase that antagonizes the autophagy-inducing kinase VPS34, thereby limiting autophagosome formation – is associated with centronuclear myopathy (Vergne *et al*., 2009). While many substrates, particularly those targeted by the UPS, have been identified, the precise determinants of substrate selection by autophagy during muscle atrophy – whether governed by receptor-mediated interactions, metabolic signaling or subcellular localization – remain incompletely understood. Given the dual role of autophagy in both preserving and degrading muscle components, a deeper understanding of its temporal dynamics and substrate specificity during atrophy is crucial.

This study provides a time-resolved, quantitative analysis of protein synthesis and degradation dynamics during TNF-α-induced atrophy in C2C12 myotubes. Using dynamic Stable Isotope Labeling with Amino Acids in Cell Culture (dynamic SILAC) (Doherty *et al*., 2009) coupled with LC-MS/MS, we analyzed the proteome landscape of early and late TNF-α-induced atrophic response while also differentiating between degraded and newly synthesized proteins. Contrary to the conventional view that atrophy is primarily driven by reduced protein synthesis and increased protein degradation, our findings reveal a more nuanced remodeling of the proteome, characterized by selective regulation of specific protein groups. Notably, while the biosynthesis of ribosomal proteins is suppressed during atrophy, mitochondrial ribosomal protein synthesis is upregulated – potentially reflecting a compensatory or adaptive response. We further explore the role of proteolytic systems, particularly autophagy, in maintaining proteostasis under atrophic conditions. Our results indicate that autophagy selectively degrades specific subsets of proteins: while myofibrillar proteins are spared, ER-resident proteins are *bona fide* autophagic cargos. We thereby uncover the potential role of ER-phagy in atrophic muscle remodeling. Understanding the contribution of the understudied autophagy pathway to skeletal muscle atrophy may uncover novel therapeutic targets for muscle-wasting disorders and bridge the knowledge gap between protein turnover regulation and metabolic adaptations in muscle atrophy.

## Results

### Dynamic SILAC identifies protein turnover changes in TNF-α-induced muscle atrophy

To investigate the details of proteostasis in skeletal muscle during homeostatic and atrophying conditions, we employed a dynamic Stable Isotope Labeling with Amino Acids in Cell Culture (dynamic SILAC) approach (Doherty *et al*., 2009) in C2C12 mouse skeletal muscle cells, both in homeostasis and TNF-α-induced atrophy (Figure 1A). Our study focuses on *de novo* protein synthesis and degradation dynamics by assessing the contributions of the autophagy-lysosome pathway (referred to as autophagy from now on) and the ubiquitin proteasome system (UPS) to these processes. To delineate their roles in protein turnover, we utilized pharmacological inhibitors prior to cell harvest – Bafilomycin A1 (BafA1) for inhibiting autophagosome-lysosome fusion and thus autophagy and Lactacystin (Lac) for inhibiting the UPS. The experimental workflow began with the differentiation of C2C12 myoblasts into mature myotubes in light-labeled medium (containing K0 = Lysine 0 [^12^C_6_, ^14^N_2_]; R0 = Arginine 0 [^12^C_6_, ^14^N_4_]). After 7 days of differentiation, myofibrillar structures are fully developed (baseline t0). At this point, the cells were switched to heavy-labeled medium (containing K8 = Lysine 8 [^13^C_6_, ^15^N_2_]; R10 = Arginine 10 [^13^C_6_, ^15^N_4_]), ensuring that every newly synthesized protein would incorporate heavy amino acids and be detected as “heavy-labeled” in mass spectrometry. This enabled us to track protein synthesis by following the accumulation of heavy-labeled peptides, while protein degradation was inferred from the disappearance of pre-existing light-labeled proteins over time. The cells were harvested after 24 hours (t24) and 72 hours (t72) to evaluate protein turnover in differentiated myotubes over a three-day period. To assess atrophic influence on proteostasis, we followed the same labelling approach but added TNF-α at the time of the medium switch (day 7) to the culture to induce muscle atrophy. Cells were again harvested 24 hours and 72 hours post-pulse. The collected cells were analyzed using liquid chromatography-tandem mass spectrometry (LC-MS/MS). We refer to 24h and 72h of TNF-α treatment as “early” and “late” time points, respectively, within the context of our experimental window. These labels are operational rather than definitive, as they are not based on a full phenotypic timeline of muscle atrophy. However, the distinct protein turnover profiles observed at these two stages suggest temporally regulated proteomic programs. These differences support the biological relevance of distinguishing between early and late responses in the atrophic process.

**Figure 1:**
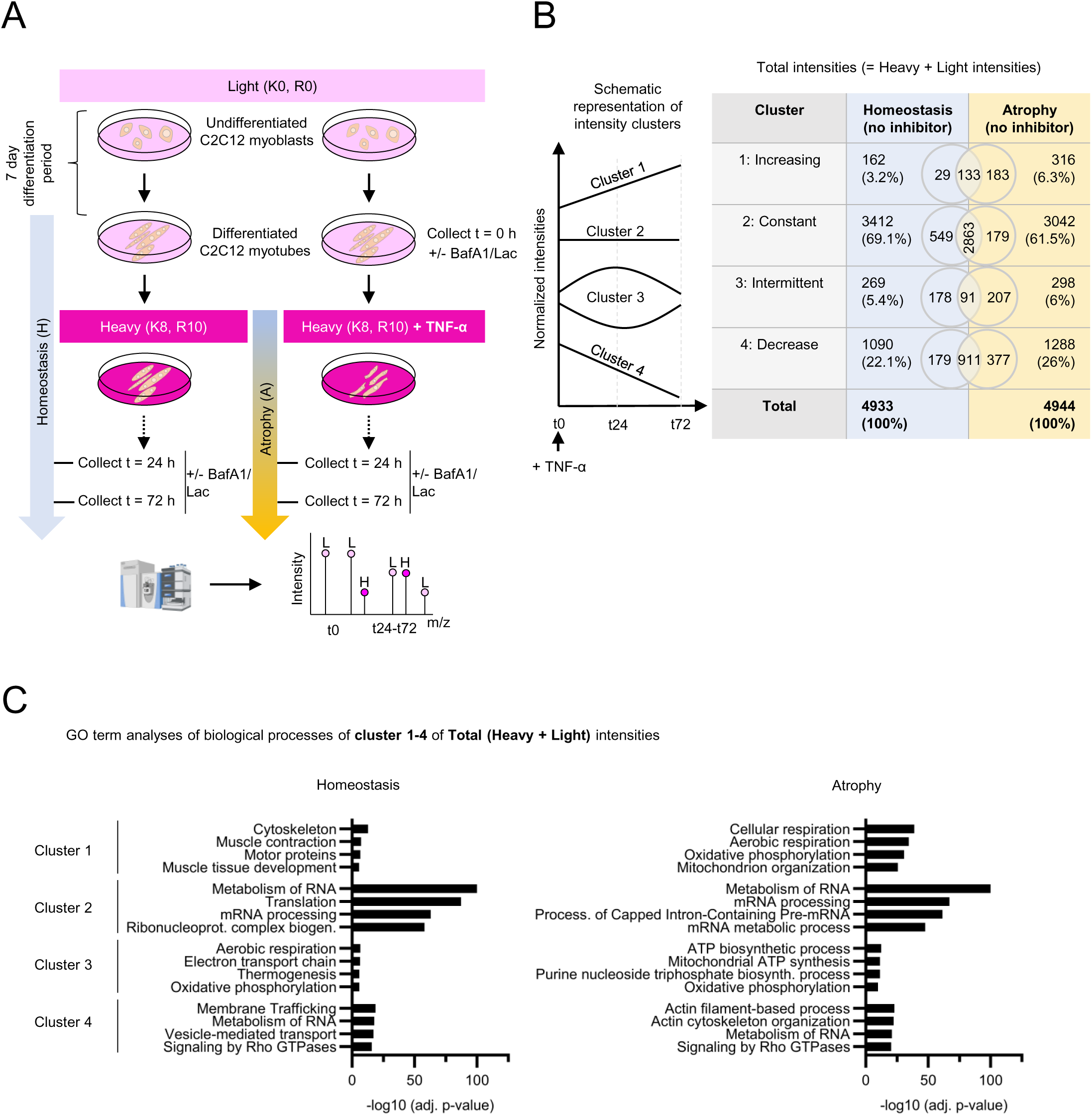
Dynamic SILAC reveals distinct sets of proteins affected in their turnover upon TNF-α-induced muscle atrophy in C2C12 cells. **(A)** Experimental workflow: C2C12 myoblasts were differentiated for seven days in light medium (K0 = Lysine 0 [^12^C_6_, ^14^N_2_]; R0 = Arginine 0 [^12^C_6_, ^14^N_4_]) into myotubes (= t0). Medium was then replaced with heavy medium (K8 = Lysine 8 [^13^C_6_, ^15^N_2_]; R10 = Arginine 10 [^13^C_6_, ^15^N_4_]) for 24 or 72 hours (t24, t72), with or without TNF-α to induce atrophy. Proteolytic inhibitors (BafA1: Bafilomycin A1 inhibiting autophagy; Lac: Lactacystin inhibiting the UPS) were administered for three hours prior to cell harvest, and cells were analyzed using LC-MS/MS. n = 3 replicates. **(B)** Protein clustering based on total intensity dynamics of non-inhibitor treated homeostatic and atrophying cells: Proteins were categorized into four clusters according to their normalized total intensity values (light + heavy) at t0, t24, and t72. Percentages of protein counts were rounded up. **(C)** Proteins of each cluster of total intensities were subjected to GO term enrichment (biological processes) analysis.

Principal component analyses (PCA) reveal distinct clustering patterns between homeostasis and atrophy samples in the heavy fraction (newly synthesized proteins) (Figure S1A-C). In contrast, in the light fractions (degraded proteins), samples treated with TNF-α for 24 hours (A24) cluster rather closely with homeostatic controls at the same time point, with separation becoming more apparent after 72 hours of atrophy (A72) (Figure S1A). Inhibition of the UPS with Lac significantly impacts the overall proteome in both heavy and light fractions, as indicated by distinct clustering across time points (Figure S1B,C). Additionally, also in both heavy and light fractions, samples treated with BafA1 and those without proteolytic inhibitor treatment cluster closely together (Figure S1C). In total, nearly 5,000 proteins were detected. Normalized light and heavy intensities were summed up to calculate the total intensities of individual proteins at time points t0, t24, and t72. Based on the temporal profiles of their total intensities, we sorted the proteins into four distinct clusters (Figure 1B): (1) those with steadily increasing intensities from t0 to t72; (2) those with constant intensities, defined as exhibiting no more than a 10% variation relative to the maximum change intensity; (3) those with intermittent intensities, characterized by an initial drop or rise at t24 followed by a return to baseline levels at t72; and (4) those with steadily decreasing intensities from t0 to t72. Interestingly, our analyses did not reveal major differences in the overall percentages of proteins with increasing or decreasing intensities between homeostatic and atrophying conditions.

However, Gene Ontology (GO) enrichment analysis (Figure 1C) of each cluster revealed distinct functional associations in atrophying and homeostatic cells, particularly in clusters 1 and 4. In homeostasis, as expected, proteins that increased their intensity over time (cluster 1) were primarily associated with the cytoskeleton, tissue development, and muscle contraction, whereas in atrophy, unexpectedly, they were linked to mitochondrial energy metabolism. Proteins summed up in cluster 4 (decreasing) in homeostasis were associated with membrane trafficking and vesicle-mediated transport while in atrophy they were predominantly related to actin filament-based process and actin cytoskeleton organization. Clusters 2 and 3 comprise similar GO terms for both, homeostasis and atrophy. While cluster 2 (constant expression) was associated with RNA metabolism and translation, cluster 3 (intermittent expression) GO terms related to oxidative phosphorylation (OXPHOS) and mitochondrial ATP synthesis. We decided to further investigate these pathways by measuring translation rates under both conditions.

### Short-term global translation inhibition in TNF-**α**-induced atrophy

To determine short-term global translation rates, we utilized the Click-iT® Plus OPP (O-propargyl-puromycin) Protein Synthesis Assay to visualize protein synthesis in both homeostatic and TNF-α-induced atrophying C2C12 myotubes (Figure 2). During OPP labeling (30 minutes), newly synthesized proteins get fluorescently tagged with a puromycin analog and appear as puncta in the cell which serve as quantification base for protein synthesis rates. Background cytosolic staining might appear due to incomplete washing, non-specific binding or excess fluorophore. As expected, treatment with the protein synthesis inhibitor cycloheximide (CHX) effectively abolished protein synthesis, validating the specificity of the OPP assay. Under homeostatic conditions (H24, H72), protein synthesis remains high over the time course. In contrast, we observed significantly lower acute protein synthesis rates in atrophying myotubes compared to homeostatic myotubes, with no significant difference in protein synthesis rates between early (A24, 24 hours) and late (A72, 72 hours) atrophy.

**Figure 2:**
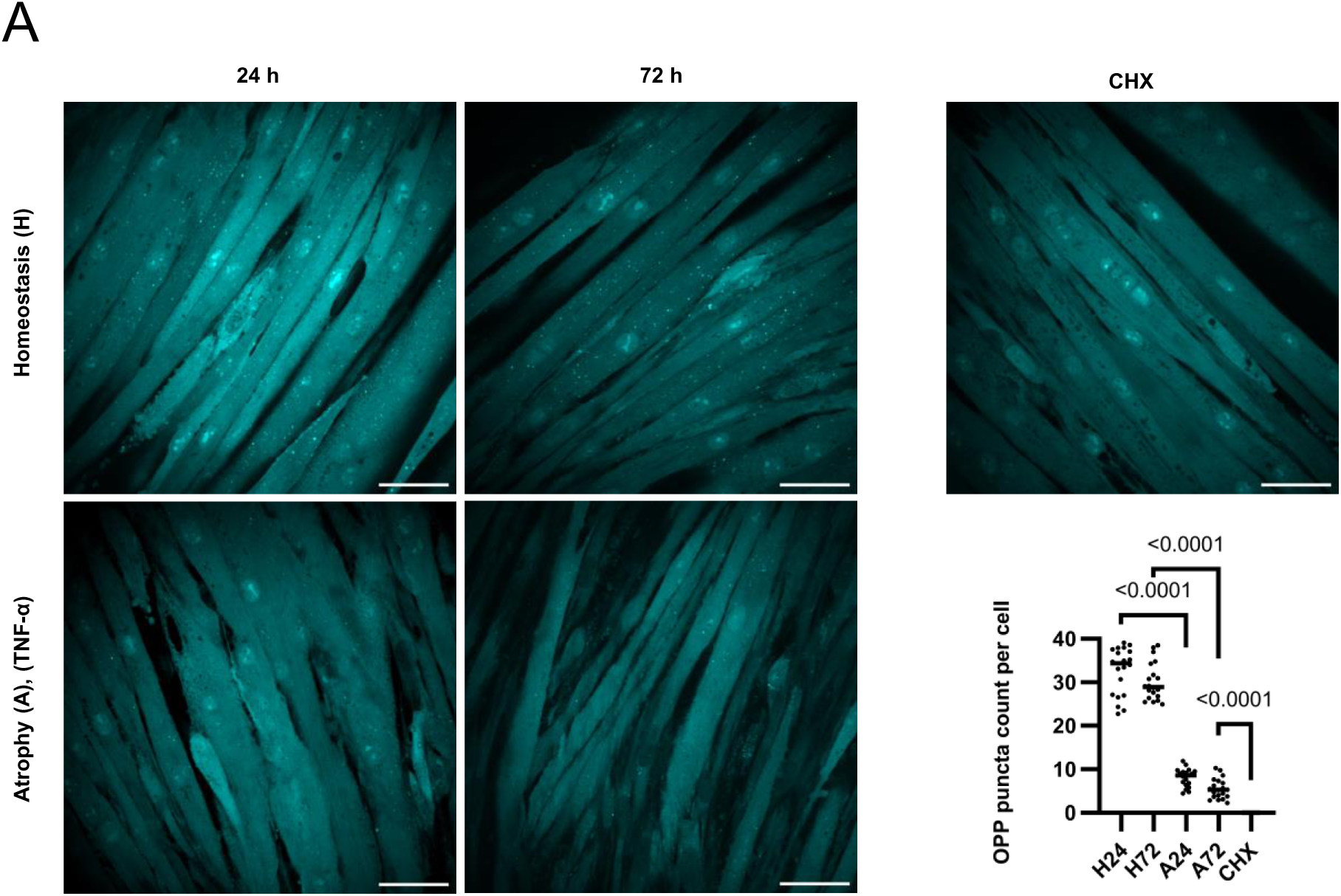
TNF-α-induced atrophy leads to acute translation inhibition. OPP (O-propargyl-puromcyin) labeling of differentiated myotubes (homeostasis at 24 h (H24) and 72 h (H27)) and TNF-α-induced atrophying C2C12 cells (24 h (A24) and 72 h (A72)) was used to measure protein synthesis rates at time points t24 and t72. Cycloheximide (CHX) was employed for 90 min as a control to inhibit protein synthesis. Puncta were manually counted across 20 images per condition, each containing over 200 cells. Statistical significance was assessed using one-way ANOVA followed by Tukey’s multiple comparisons test. Scale bar: 60 µm.

### Dynamic SILAC analysis identifies early and late-phase protein degradation and synthesis signatures during TNF-**α**-induced atrophy

Proteome analysis via dynamic SILAC enables the distinction between newly synthesized proteins (Heavy channel) and pre-existing proteins (Light channel), eliminating reliance on total intensity measurements. To assess protein turnover over time in both atrophic and homeostatic conditions, we performed differential abundance analysis using limma (Ritchie *et al*., 2015) with moderated t-tests and ANOVA (empirical Bayes), applying a p ≤ 0.05 threshold and Benjamini-Hochberg correction. We compared the abundance of pre-existing proteins (Light channel) at 72 hours (t72) relative to 24 hours (t24). Proteins showing significant reduction (log_2_FC < 0, adj. p-value < 0.05) were classified as actively degraded. Similarly, newly synthesized proteins were identified by comparing the Heavy channel at t72 to t24, with significant accumulation (log_2_FC > 0, adj. p-value < 0.05) indicating active synthesis (Figure 3A). Interestingly, we observed no major changes in overall numbers of newly synthesized or degraded proteins during two days of atrophy (A72 vs. A24) compared to two days of homeostasis (H72 vs. H24). Therefore, we decided to perform GO analyses to determine differentially regulated proteins: During homeostasis (H72 vs. H24), we identified 1,197 newly synthesized proteins, primarily enriched in GO terms related to mitochondria-dependent energy metabolism but also cytoskeleton, and 1,263 degraded proteins, associated with RNA metabolism, membrane trafficking, and protein transport. In contrast to that, within two days of atrophy (A72 vs. A24) 1,229 proteins were newly synthesized and 1,409 were degraded. GO analysis revealed that proteins significantly synthesized during two days of atrophy were also involved in mitochondrial energy production, predominantly OXPHOS), but interestingly, also proteins involved in processing in the ER, and ER organization were increasingly synthesized. In contrast to homeostatic conditions, degraded proteins during atrophy were associated with translation, protein transport and folding, actin filament-based processes, cytoskeletal organization, proteasome assembly, ubiquitin-mediated proteolysis, and motor proteins. To sum up, despite similar amounts of overall proteins affected by protein turnover, we could clearly observe a distinctly regulated proteome landscape with atrophy-specific synthesis of proteins enriched in ER organization and protein processing. Degraded proteins during atrophy were related to translation, cytoskeletal structure, protein folding and proteasome-related pathways, indicating a reprogramming of cellular organization and maintenance mechanisms under prolonged atrophic stimulus.

**Figure 3:**
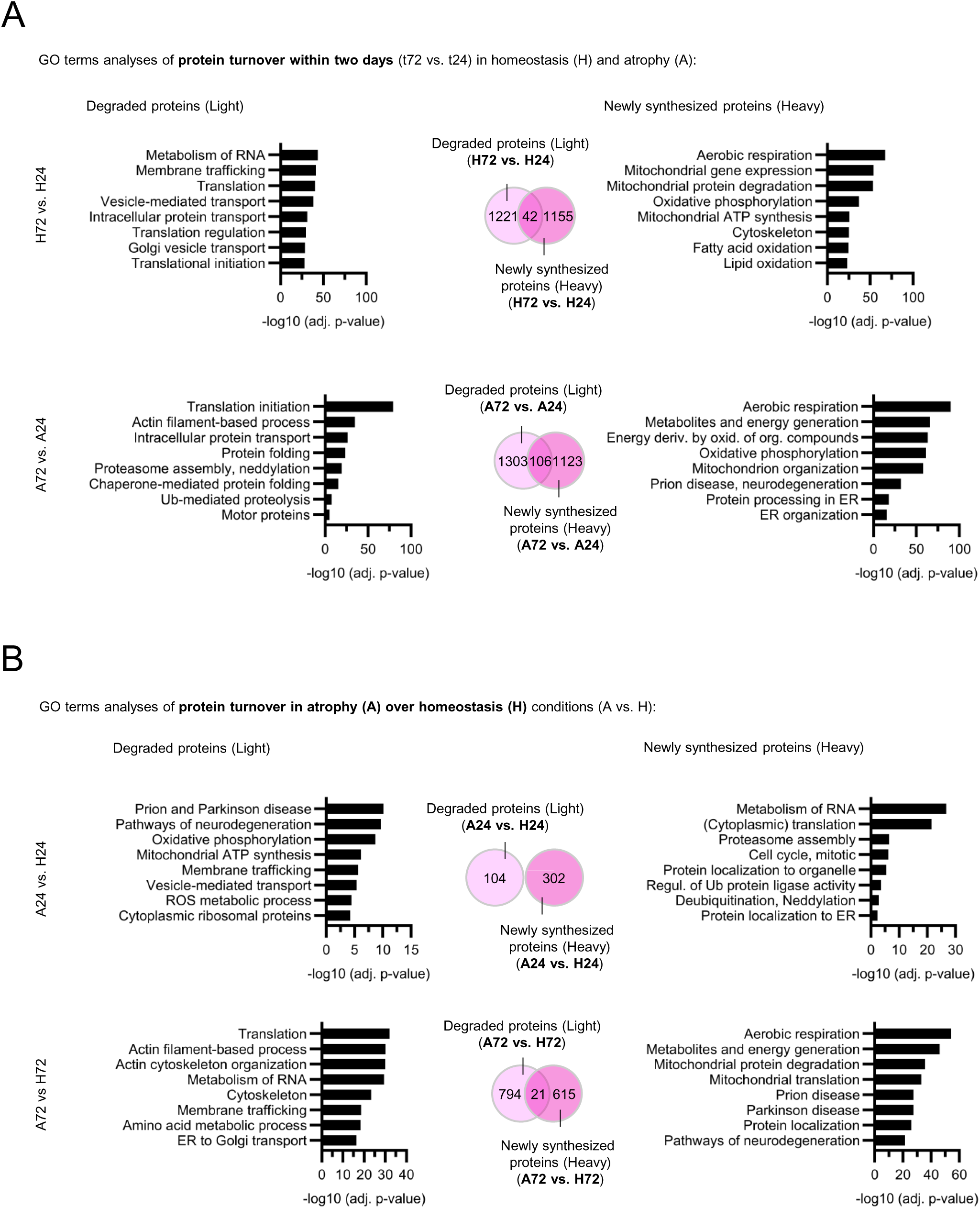
Dynamic SILAC analysis identifies early and late-phase protein degradation and synthesis signatures during TNF-α-induced atrophy. Stepwise ANOVA comparisons were performed on heavy and light dynamic SILAC samples to assess significant protein degradation and synthesis rates across and between homeostatic (H) and atrophic (A) conditions. Proteins with log_2_FC < 0 and adj. p-value < 0.05 in the light channel were classified as significantly degraded, while proteins with log_2_FC > 0 and adj. p-value < 0.05 in the heavy channel were classified as significantly newly synthesized. Numbers depict total amounts of significantly regulated proteins in respective dataset comparison. GO term analyses (biological processes) of **(A)** protein turnover within two days (t72 vs. t24) in homeostasis and atrophy and **(B)** protein turnover in atrophy over homeostasis (A vs. H). Numbers in Venn diagrams depict amount of significantly regulated proteins in respective dataset comparison. Light pink resembles light channel, dark pink resembles heavy channel

To distinguish early and late atrophic responses relative to homeostasis, we also compared A24 vs. H24 and A72 vs. H72 (Figure 3B). Again, proteins showing significant reduction in the light channel (log_2_FC < 0, adj. p-value < 0.05) were classified as actively degraded, whereas newly synthesized proteins were identified with log_2_FC > 0 and an adj. p-value of < 0.05. In early TNF-α response (A24 vs. H24), 302 newly synthesized proteins specific to atrophy were detected, with no overlap with the 104 degraded proteins identified. GO analysis of degraded proteins in A24 vs. H24 highlighted pathways associated with protein misfolding and aggregation (neurodegeneration-related GOs), ATP synthesis, protein transport, ROS metabolic processes, and cytoplasmic ribosomal protein degradation. Newly synthesized proteins in A24 vs. H24 were linked to cytoplasmic translation, cell cycle regulation, UPS components, and protein localization to the ER. In late atrophy (A72 vs. H72), the number of newly synthesized proteins specific to atrophy nearly doubled (626) and degraded proteins increased to 815. GO analyses of degraded proteins in late TNF-α response depicted pathways related to translation, actin cytoskeleton, membrane trafficking and amino acid metabolism whereas newly synthesized proteins were related to cellular energy metabolism, degradation and translation of mitochondria, neurodegeneration-related GOs and protein localization. To sum up, comparison of the early and late TNF-α-induced atrophic proteome to the proteome of homeostatic cells revealed a marked increase in both newly synthesized and actively degraded proteins, indicating a shift in the cellular program from 24 hours to 72 hours of TNF-α exposure toward enhanced protein turnover, mitochondrial remodeling and stress adaptation.

### Selective degradation and impaired synthesis of myofibrillar proteins during TNF-**α**-induced muscle atrophy

Skeletal muscle atrophy is characterized by the breakdown of myofibrillar components which we could also observe in our previous GO analyses (Figure 3B, “actin-filament based processes”, “motor proteins”). However, an in-depth analysis focusing on the specific proteins targeted for degradation during different atrophy stimuli remains lacking. To address this gap, we categorized myofibrillar proteins into the previously defined clusters (Figure 1B) of total intensities to assess their absolute dynamics under homeostatic and atrophic conditions (Figure S2). Generally, we could observe that the majority of myofibrillar components show stable intensities from t0 to t72 in both conditions (therefore assigned to cluster 2). In both conditions, the majority of myofibrillar components was assigned to cluster 2, indicating stable intensities from t0 to t72. However, in atrophy, as expected, a greater proportion of myofibrillar proteins were sorted into cluster 4 (decreasing intensities from t0 to t72), reflecting overall decrease. Conversely, fewer myofibrillar proteins were assigned to cluster 1 (increasing intensities from t0 to t72) in atrophy. Notably, Myh4, one of the myosin heavy chain proteins which acts as a critical component to sustain contraction, was the only myofibrillar protein that was assigned to cluster 1 under atrophic conditions but was classified in cluster 2 (constant intensities) under homeostasis. These findings prompted us to further examine degradation and synthesis of individual myofibrillar components in TNF-α-induced atrophying conditions and how these changes compare to homeostatic conditions (Figure 4) instead of observing total intensities (Figure S2). Proteins in close proximity to actin filaments (Actn1-4), including Capza1,2 and Flna-c but also proteins located at the M band (Fhl1-3, Myl, Tpm2-4) exhibited pronounced degradation in atrophic conditions (A72 vs. H72) (Figure 4), whereas myofibrillar protein degradation was not abolished in homeostatic conditions (H72 vs. H24) due to normal protein turnover. Myofibrillar protein synthesis in comparisons of atrophy vs. homeostasis (A24 vs. H24 and A72 vs. H72) showed no significant upregulation, whereas, in stark contrast, homeostatic conditions (H72 vs. H24) demonstrated high synthesis rates of myofibrillar proteins. However, myofibrillar protein synthesis was not entirely abolished in atrophy. The comparison between late and early atrophy (A72 vs. A24) reflecting synthesis over two days of atrophy, revealed some synthesis that was downregulated compared to two days of homeostasis, with mostly myosin heavy chain (Myh) proteins again being newly synthesized in late-stage atrophy. Interestingly, all monitored myofibrillar proteins were either newly synthesized or degraded, but not both, suggesting inverse regulation of protein turnover at the single-protein level. In summary, in-depth analysis of differentially degraded and newly synthesized myofibrillar proteins shows increased degradation in late-stage atrophy and moderate but lower degradation in homeostatic conditions compared to atrophying conditions. In contrast, myofibrillar protein synthesis is reduced in atrophy compared to homeostasis.

**Figure 4:**
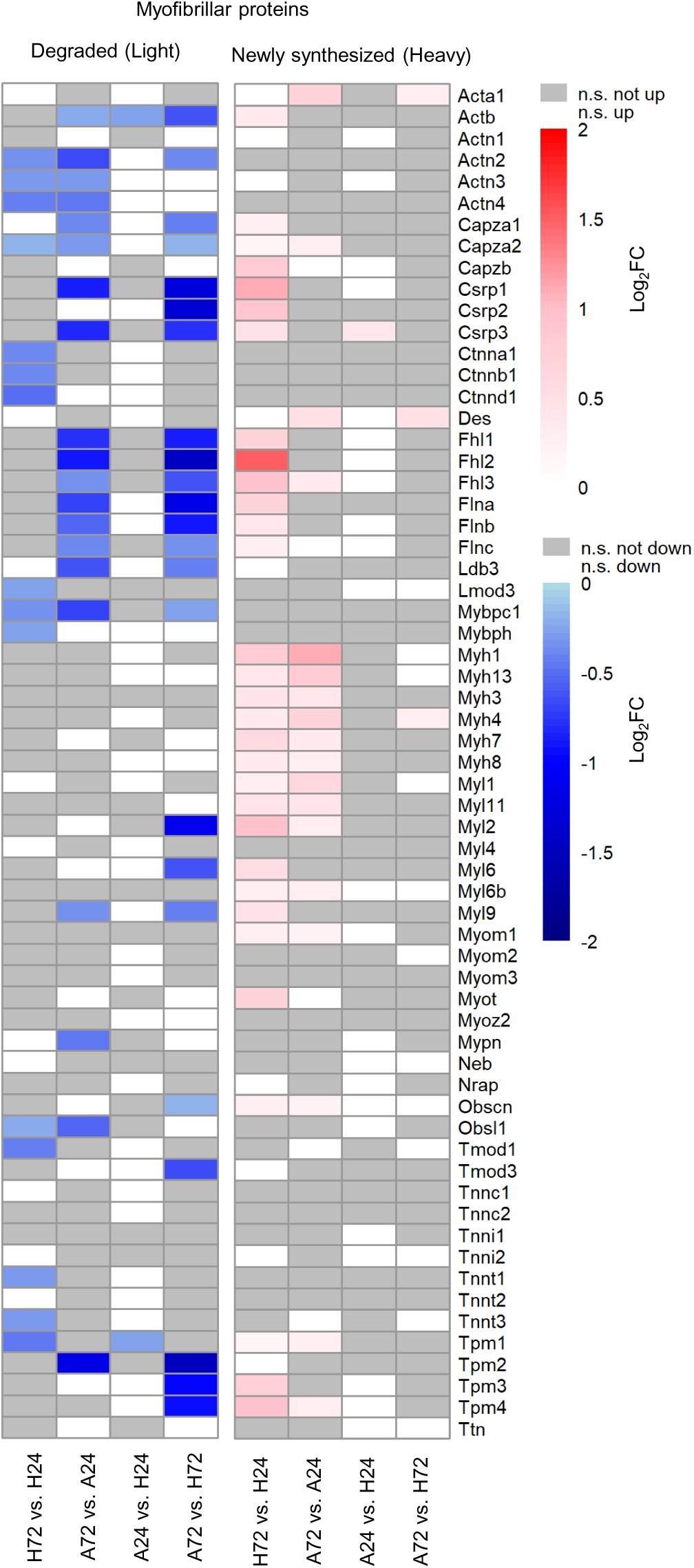
Elevated degradation and reduced synthesis of myofibrillar components upon TNF-α-induced atrophy. Stepwise ANOVA comparisons were performed on heavy and light dynamic SILAC samples to assess significant protein degradation and synthesis rates across and between homeostatic (H) and atrophic (A) conditions. Proteins with log_2_FC < 0 and adj. p-value < 0.05 in the light channel were classified as significantly degraded, while proteins with log_2_FC > 0 and adj. p-value < 0.05 in the heavy channel were classified as significantly newly synthesized. Heatmap of degraded and newly synthesized myofibrillar proteins. White: not significantly up/downregulated; grey: not significant.

### Reduced synthesis of cytoplasmic ribosomal proteins contrasts with increased mitochondrial ribosomal protein synthesis during muscle atrophy

Ribosomal proteins underwent pronounced degradation during two days of atrophy (A72 vs. A24), exceeding degradation observed during the equivalent homeostatic period (H72 vs. H24) (Figure 5A), reflecting reduced translation rates (Figure 2). Moreover, degradation rates of ribosomal proteins were notably higher in late atrophy compared to homeostasis (A72 vs. H72) than in early atrophy (A24 vs. H24). Interestingly, ribosomal protein synthesis occurred substantially during the two-day homeostatic period (H72 vs. H24) and persisted into early atrophy (A24 vs. H24). However, synthesis of ribosomes was nearly absent as evidenced by comparisons of late stage atrophy with homeostasis (A72 vs. H72) and early atrophy (A24 vs. H24). Given the prevalent mitochondrial energy metabolism-related GO terms identified previously, we further investigated mitochondrial ribosomal proteins in detail (Figure 5B). We observed a modest increase in degradation under atrophic conditions relative to homeostasis (A24 vs. H24 and A72 vs. H72). Interestingly, mitochondrial ribosomal proteins displayed reduced synthesis rates in early atrophy compared to homeostasis, but synthesis notably increased over two days of atrophy (A72 vs. A24), surpassing the synthesis levels observed during two days of homeostasis (H72 vs. H24). Similar to myofibrillar proteins, ribosomal and mitochondrial ribosomal proteins exhibited an inverse pattern of regulation with individual proteins undergoing either degradation or synthesis, but not both, across the examined conditions.

**Figure 5:**
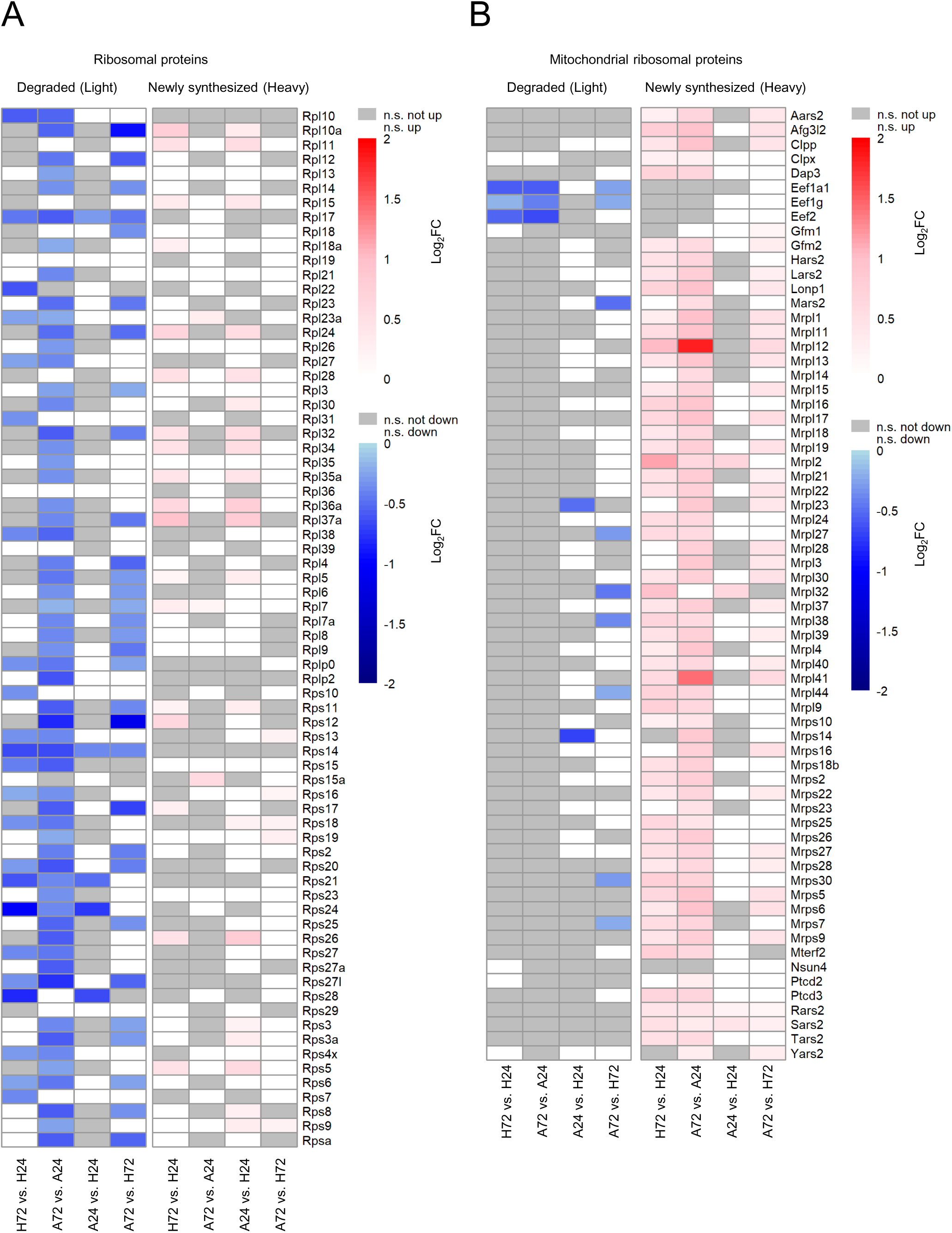
Differential regulation of ribosomal and mitochondrial ribosomal proteins upon TNF-α -induced atrophy. Stepwise ANOVA comparisons were performed on heavy and light dynamic SILAC samples to assess significant protein degradation and synthesis rates across and between homeostatic (H) and atrophic (A) conditions. Proteins with log_2_FC < 0 and adj. p-value < 0.05 in the light channel were classified as significantly degraded, while proteins with log_2_FC > 0 and adj. p-value < 0.05 in the heavy channel were classified as significantly newly synthesized. Heatmap of degraded and newly synthesized ribosomal **(A)** and mitochondrial ribosomal **(B)** proteins. White: not significantly up/downregulated; grey: not significant.

### Increased autophagic turnover upon TNF-**α**-induced atrophy assessed by LC3-II flow cytometry

To assess autophagic activity under atrophying conditions, we quantified LC3-II levels in C2C12 cells using a flow cytometry-based approach (Alsaleh *et al*., 2020). This approach builds on previous flow cytometry studies in C2C12 myotubes (Nolan *et al*., 2024), extending it to monitor LC3-II dynamics and is more quantitative than Western blot (Alsaleh *et al*., 2020). C2C12 cells were differentiated for 7 days, followed by 24 hours or 72 hours incubation with or without TNF-α to model homeostatic and atrophying conditions. To evaluate autophagic activity, cells were treated for 3 hours with BafA1 to stop autophagosome-lysosome fusion, or vehicle-treated (DMSO) before harvest. BafA1 causes accumulation of LC3B-II positive autophagosomes that would otherwise be degraded, resulting in a shift in the LC3-II signal (Figure 6A). LC3B-I is washed out with a mild detergent treatment before staining intracellularly for LC3B-II only. The gating strategy is shown in Supplementary Figure 3. Interestingly, LC3B-II signal intensity was mildly but significantly increased among DMSO-treated samples under atrophying conditions compared to homeostasis (Figure 6B), suggesting increased basal autophagosome synthesis. This was accompanied by increased autophagic turnover at 24 hours of atrophy (Figure 6C, H24 vs. A24), quantified as the difference in LC3B-II signal between BafA1-and DMSO-treated samples similar to 72 hours of atrophy, although not significant (Figure 6C, H72 vs. A72). These findings suggest an increase of autophagy in atrophy.

**Figure 6:**
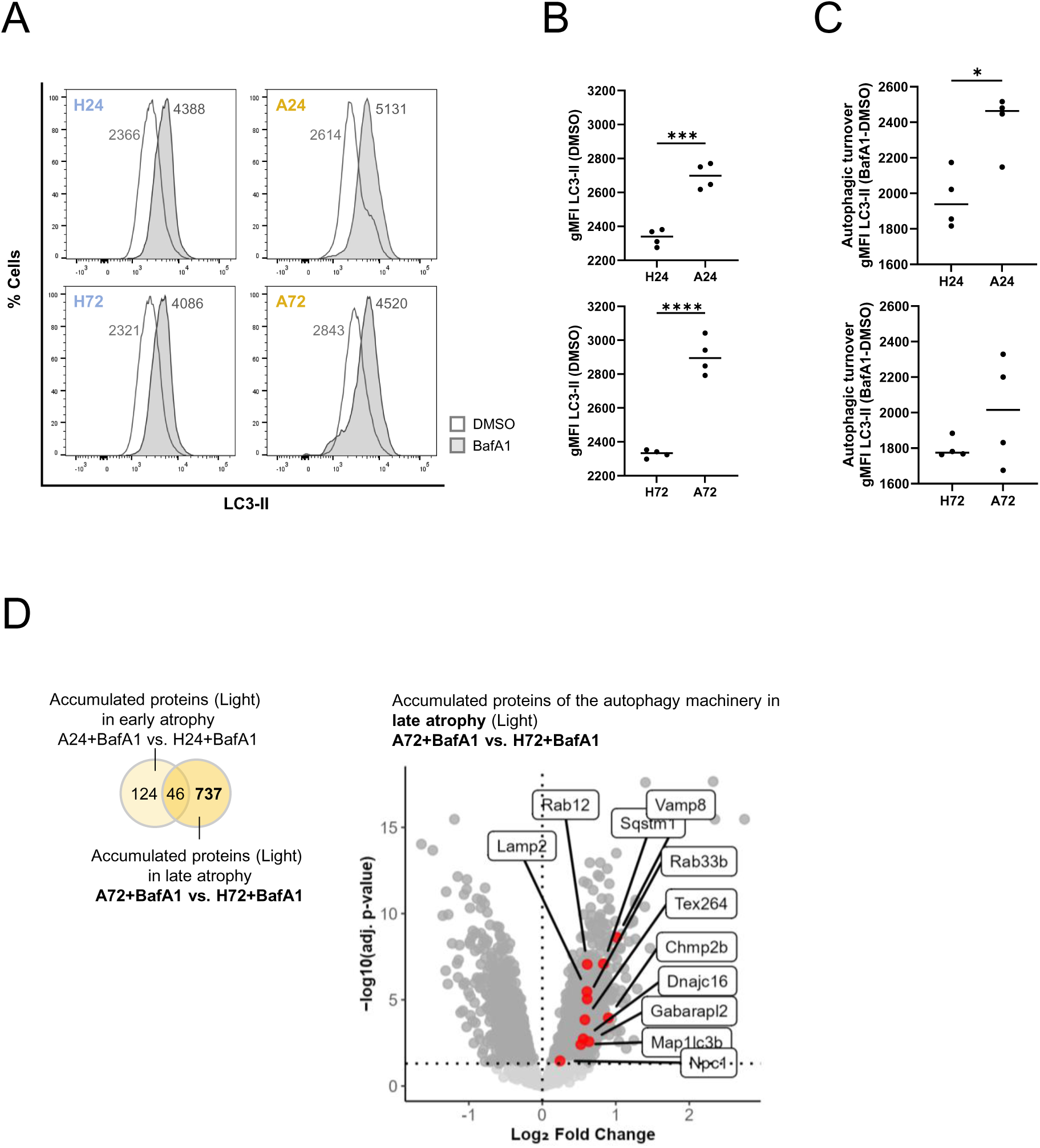
Increased autophagic turnover in TNF-α-induced atrophying C2C12 cells. **(A-C)** Flow cytometry-based assay measuring autophagic turnover by staining membrane-bound LC3-II with a fluorescently labeled antibody (Alsaleh *et al*., 2020) **(A)** Representative histograms display the shift in LC3-II signal following 3 h BafilomycinA1 (BafA1) treatment **(B)** Elevated LC3-II signal intensity (gMFI, geometric mean fluorescence intensity) in atrophying (A24, A72) cells compared to homeostatic cells (H24, H72) in dimethylsulfoxide (DMSO)-treated samples. N = 4 replicates. **(C)** Autophagic turnover was calculated by determining the difference in LC3-II signal intensity (gMFI) between BafA1-treated and DMSO-treated cells over a 3h incubation period. N = 4 replicates. **(D)** Accumulated proteins upon BafilomycinA1 (BafA1) in ANOVA comparisons of atrophying (A24+BafA1 and A72+BafA1) vs homeostatic (H24+BafA1 and H72+BafA1) C2C12 cells in the light channel with log_2_FC set > 0 and adj. p-value set at < 0.05. Numbers in Venn diagram depict amount of significantly regulated proteins in respective dataset comparison. Red dots shown in volcano plot of A72+BafA1 vs. H72+BafA1 depict annotated autophagic machinery components.

To validate this, we analyzed the accumulation of autophagy machinery components in BafA1-treated atrophying conditions compared to BafA1-treated homeostatic conditions in our dynamic SILAC dataset (Figure 6D). Proteins with a significant increase in the light channel (log_2_FC > 0, adj. p-value < 0.05) were considered to accumulate as a result of inhibited autophagic turnover. Strikingly, a markedly higher number of proteins accumulated at 72 hours of atrophy compared to 24 hours of atrophy with 783 and 170 significantly enriched proteins, respectively, and minimal overlap (n = 46). In line with our flow cytometry–based analysis, Map1lc3b was among the proteins that accumulates. Additionally, other autophagy-related factors – including Lamp2, Tex264, Rab33b, and Gabarapl2 – accumulated as well.

### Autophagic receptor p62/Sqstm1 is localized at the M band of the sarcomere but lacks additional autophagic machinery

To gain further insight into the function of autophagy in skeletal muscle (particularly during atrophy), we aimed to locate the autophagy machinery within the sarcomere. The protein p62/Sqstm1 functions as a selective autophagy receptor, enabling it to specifically target ubiquitylated substrates for engulfment by autophagosomes, which are subsequently degraded upon fusion with lysosomes. In immunocytochemical stainings of (atrophying) C2C12 cells, we observed p62/Sqstm1 puncta formation, co-localized with myosin heavy chain (MyHC), quantified with line plots along the sarcomere (Figure 7A-B). This is in line with observations in cardiomyocytes (Lange *et al*., 2005; McNamara *et al*., 2023) and indicates p62/Sqstm1 localization at the M band of the sarcomere in the skeletal muscle. However, we did not observe any strong colocalization of other autophagy machinery components like LC3B, Wipi2 (phagophore marker), lysin 63-linked ubiquitin chains (K63-Ub) and Lamp1 (lysosomal marker) with MyHC in neither C2C12 myotubes (Figure S4) nor TNF-α-induced atrophying myotubes (Figure S5), suggesting a lack of accumulation or functional assembly of the broader autophagy machinery in this specific region of the sarcomere.

**Figure 7:**
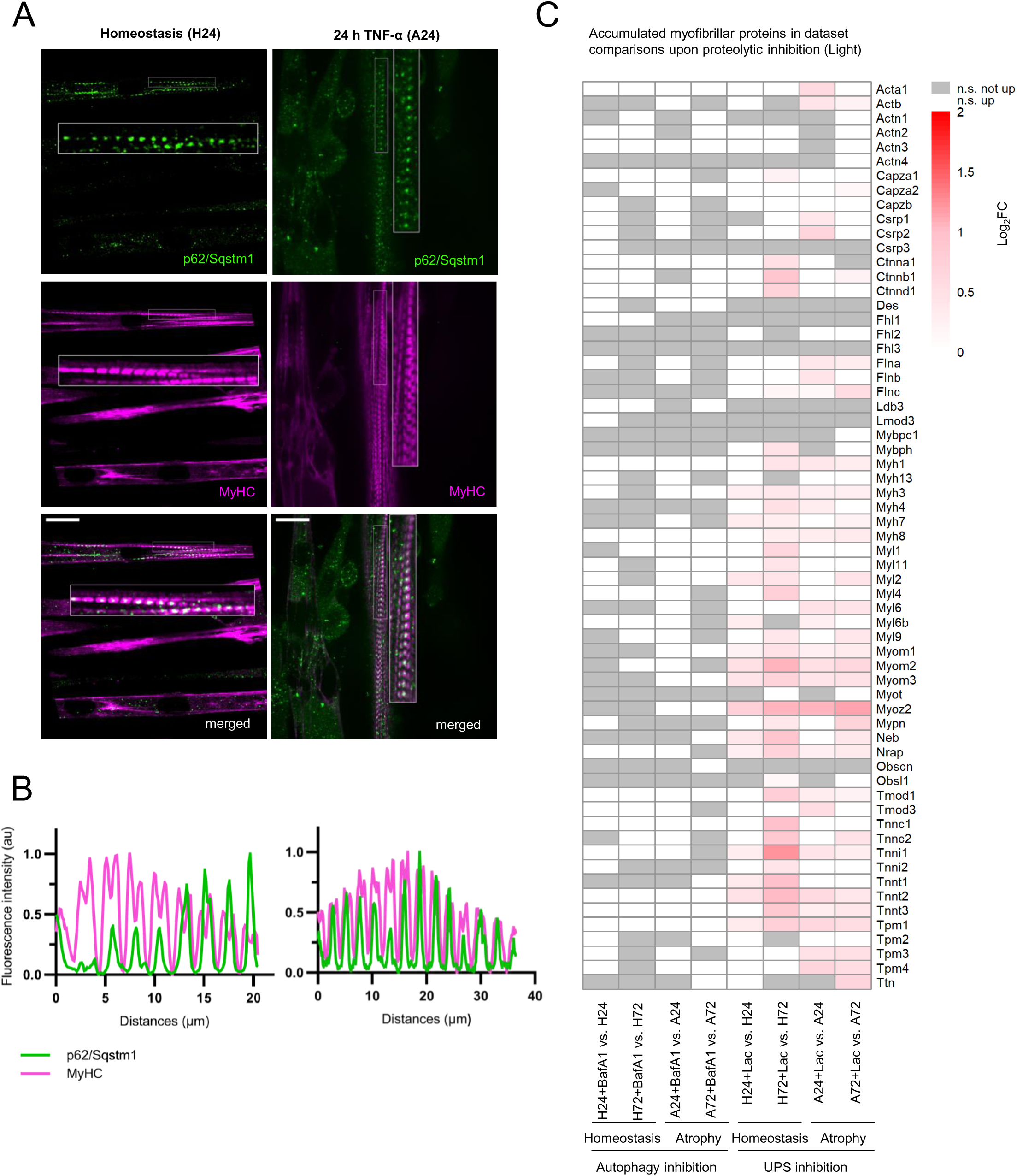
Autophagy machinery is not involved in myofibrillar protein degradation. **(A)** Immunocytochemistry staining of C2C12 (atrophying) myotubes for p62/Sqstm1 and myosin heavy chain (MyHC) highlighting the M band region of the sarcomere. **(B)** Corresponding line plots display fluorescence intensity profiles for each channel along the length of the sarcomere. Scale bar 20 µm. **(C)** Accumulated myofibrillar proteins upon BafilomycinA1 (BafA1) and Lactacystin (Lac) treatment in ANOVA comparisons of homeostatic (H) and atrophying (A) C2C12 cells in the light channel with log_2_FC set at > 0 and adj. p-value set at < 0.05. White: not significantly accumulated; grey: not significant.

### Inhibition of autophagy does not cause accumulation of myofibrillar proteins

Given the presence of p62/Sqstm1 at the M band in the absence of other autophagy machinery components, we investigated whether autophagy contributes to the degradation of myofibrillar proteins. To assess the influence of either proteolytic system, i.e. autophagy and UPS, on the degradation of myofibrillar proteins, proteolytically inhibited and non-inhibited samples were compared using the pre-existing protein pool (Light channel) (Figure 7C). Proteins showing significant increase in the light channel (log_2_FC > 0, adj. p-value < 0.05) were classified as accumulated upon inhibition. Inhibition of the UPS (Lac) led to significant accumulation of numerous myofibrillar proteins from all regions of the sarcomere, in both homeostatic and atrophying myotubes, with the strongest accumulation observed in homeostatic conditions (H72+Lac vs. H72). Conversely, inhibition of autophagy (BafA1) did not result in significant accumulation of myofibrillar proteins, suggesting minimal or no involvement of autophagy in their degradation.

### Shift in autophagic selectivity from homeostatic ER and metabolic maintenance to ER-stress-linked degradation in TNF-**α**-induced atrophy

To further characterize the autophagic cargo selectively targeted during atrophy compared to homeostasis, we performed GO analyses on proteins accumulating upon autophagy inhibition in both conditions (Figure S6A, S8C). Again, proteins showing significant increase in the light channel (log_2_FC > 0, adj. p-value < 0.05) were classified as accumulated upon autophagic inhibition. Under homeostatic conditions (H+BafA1 vs. H), 73 proteins accumulated compared to 91 accumulating in atrophic conditions (A+BafA1 vs A) upon autophagy inhibition with an overlap of 38 proteins in both conditions. Both homeostasis and atrophy share a core set of enriched terms upon inhibition, including ECM-related processes, cytoskeletal components, and ER-localized multiprotein complexes. However, proteins accumulating in atrophy were additionally associated with protein folding, maturation, and ER stress response (like Hspa5/BiP) (Figure S6A, S7, S8, Figure 8B), whereas proteins accumulating in homeostasis were involved in metabolic processes such as alcohol and cholesterol metabolism and cellular response to nitrogen compounds (Figure S6A, S7A, S8A-C). Furthermore, upon autophagy inhibition in atrophic conditions, more BCAA transporter were found to be accumulated than in homeostatic conditions (e.g. Slc3a2 in early and Slc7a5/Lat1 in late atrophy (Figure S7A, S8A-B). To conclude, whilst autophagy inhibition in homeostatic conditions led to accumulation of proteins related to metabolic processes and ER quality control, autophagy inhibition in atrophying condition led to accumulation of proteins related to ER quality control as well as ER stress – indicating a shift in autophagic selectivity during atrophy.

**Figure 8:**
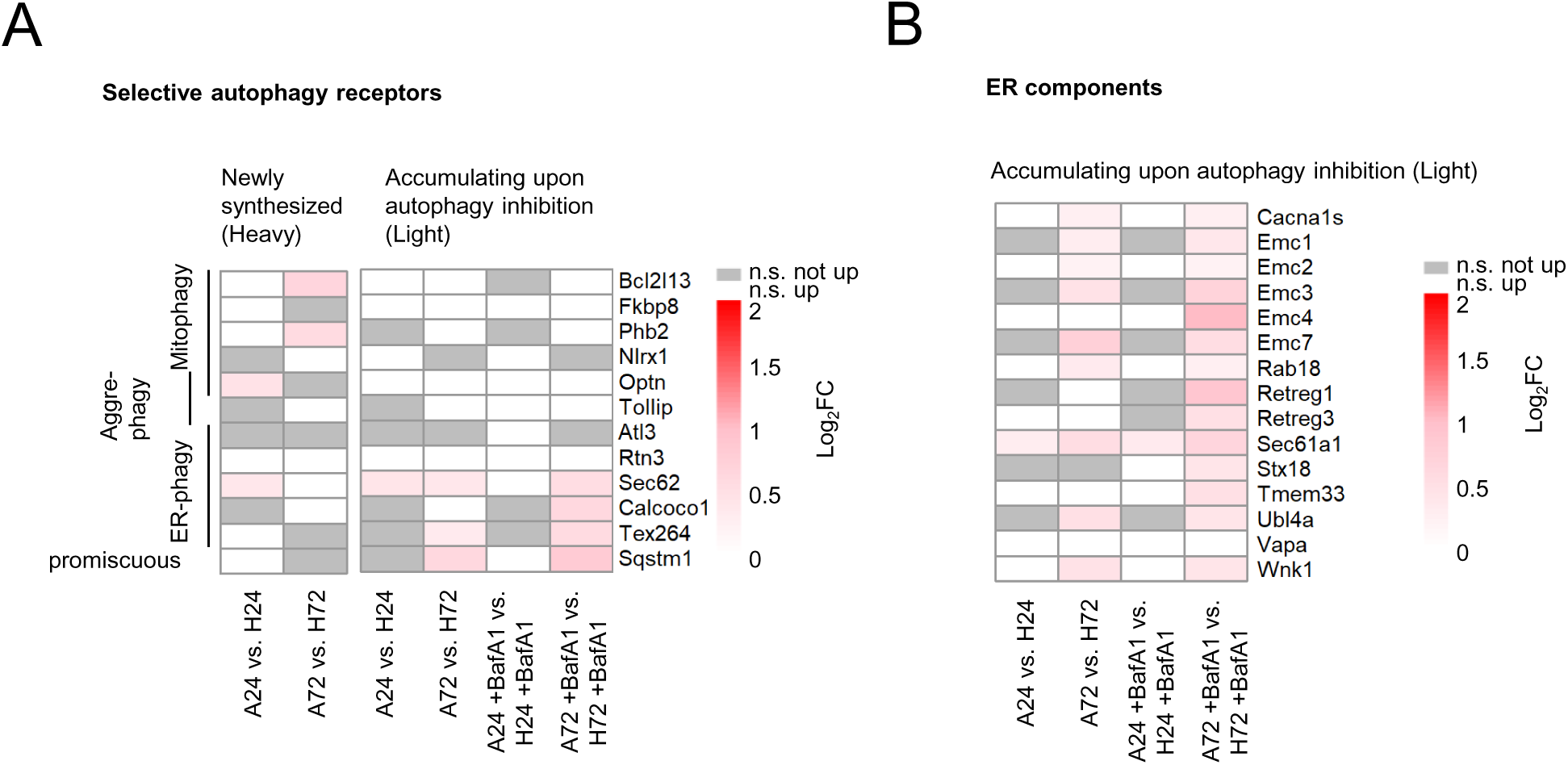
ER-phagy is the most prominent selective autophagy type during TNF-α-induced atrophy. **(A)** Left: Newly synthesized (Heavy channel) in atrophy (A24 vs. H24 and A72 vs. H72) in ANOVA comparisons with log_2_FC > 0 and adj. p-value set at < 0.05. Right: Accumulating (Light channel) selective autophagy receptors with and without BafilomycinA1 (BafA1) treatment in atrophy (A24 vs. H24 and A72 vs. H72) with log_2_FC > 0 adj. p-value set at < 0.05. **(B)** Accumulating (Light channel) ER components with and without BafilomycinA1 (BafA1) treatment in atrophy (A24 vs. H24 and A72 vs. H24) in ANOVA comparisons with log_2_FC > 0 and adj. p-value set at < 0.05. White: not significantly accumulated; grey: not significant.

To investigate ER-phagy in atrophy progression in muscle, we first established a baseline of autophagic selectivity under physiological conditions, by examining proteins accumulating under homeostatic conditions at early (H24+BafA1 vs. H24) and late (H72+BafA1 vs. H72) time points (Figure S8A,B), and assessed their enrichment for specific Gene Ontology (GO) terms (Figure S8C). Again, proteins showing significant increase in the light channel (log_2_FC > 0, adj. p-value < 0.05) were classified as significantly accumulated. At 24 hours, enriched GO terms were linked to ECM remodeling, cell migration/motility, developmental pathways and inflammatory/metabolic responses. By 72 hours, GO terms comprise pathways like membrane trafficking, cell-cell communication and vascular/neural remodeling. With this baseline established, we next distinguished early and late atrophic conditions by analyzing GO terms associated with proteins accumulating upon BafA1 treatment in early (A24+BafA1 vs. A24) and late atrophy (A72+BafA1 vs. A72) (Figure S7B, S8A,B). A total of 65 proteins accumulated in early atrophy, while 56 proteins accumulated in late atrophy, with 30 shared hits between both conditions. Both early and late atrophy showed a strong enrichment for extracellular matrix (ECM)-related processes (e.g. Col1a2, Fn1), ECM-receptor interaction, vasculature development and cytoskeleton-related processes. Similarly, amyloid-beta clearance (e.g. Lrp1, Lrp4) and regulation of nervous system development (e.g. Ephb3, App) were enriched in both early and late atrophy. Importantly, proteins related to protein folding (e.g. Pdia4, Hsp90b1) and ER-resident proteins (e.g. Tor1a, Nomo1) were accumulated in both states. Nevertheless, the proteome accumulating with BafA1 during early atrophy showed more hits related to an ER stress response (e.g. Pdia3, Hspa5, Dnajc3), the ERAD pathway and the UPR (proteasome subunits). In addition, BafA1 treatment in late atrophy led to accumulation of more autophagosome related hits compared to early atrophy (Figure S9): Gabarapl2, Map1lc3a, Stx12, Rab11b and also to a higher accumulation of autophagy receptors Calcoco1 and p62/Sqstm1, indicating increased abundance of autophagosomes as it is expected after BafA1 treatment and elevated autophagic activity in atrophy. These findings are consistent with an emphasis on ER-phagy during atrophy.

### ER-phagy is the most prominent selective autophagy type during TNF-**α**-induced atrophy

To further elucidate the specificity of autophagic response in muscle atrophy, we investigated which selective autophagy pathways are engaged or disrupted during early and late stages, respectively (Figure 8A). Proteins showing significant increase in the heavy channel (log_2_FC > 0, adj. p-value < 0.05) were classified as significantly newly synthesized. Among all published selective autophagy receptors (reviewed by Gubas and Dikic, 2022), we observed minimal new synthesis of selective autophagy receptors comparing early atrophy to homeostasis (A24 vs. H24), only Optn (mitophagy/aggrephagy) and Sec62 (ER-phagy) increased indicating a response to unfolded proteins and ER stress (Fig 8A left). Interestingly, in late-stage atrophy (A72 vs. H72), two mitophagy receptors (Bcl2l13 and Phb2) were newly synthesized but no ER-phagy receptors were synthesized (Figure 8A left). Autophagy receptors are known to accumulate upon BafA1 treatment due to impaired lysosomal degradation. To further explore this, we examined which receptors accumulate significantly in atrophying conditions compared to homeostatic conditions upon autophagy inhibition (light channel, log_2_FC > 0, adj. p-value < 0.05) (Figure 8A right). Interestingly, in early atrophy, none of the monitored autophagy receptors accumulated, suggesting functional autophagic turnover. In contrast, we detected an accumulation of three ER-phagy receptors, Sec62, Calcoco1 and Tex264, as well as p62/Sqstm1 with or without BafA1 treatment in atrophy. To further investigate ER-phagy dynamics, we monitored ER-resident proteins and found that while early atrophy showed negligible accumulation, late atrophy resulted in marked accumulation of ER components (Figure 8B). Collectively, these data indicate that autophagy is degrading ER, reaching its peak at 72 hours of atrophy.

## Discussion

This study presents a direct, time-resolved analysis of protein turnover in TNF-α-induced muscle atrophy via dynamic SILAC (Doherty *et al*., 2009), refining the conventional view that atrophy is always accompanied by decreased protein synthesis and increased protein degradation (Sandri, 2013; Bodine and Baehr, 2014). Instead, our findings reveal a selective remodeling of protein dynamics, with distinct protein groups undergoing differential regulation, indicative of an adaptive rather than purely degenerative process (as suggested by Baehr *et al*., 2017). This aligns with previous studies showing that different atrophic and hypertrophic stimuli elicit varied responses in synthesis and degradation, such as denervation of the rat diaphragm, which leads to both increased protein degradation and enhanced protein synthesis (Argadine *et al*., 2009). We demonstrate a dynamic shift in the autophagic response, transitioning from a rather homeostasis-maintaining process to elevated autophagic turnover rates with ER stress-driven mechanisms during atrophy.

We employed OPP labeling and dynamic SILAC to provide a snapshot of short-term translation rates (30 minutes) and long-term protein turnover dynamics respectively. The combination of these methods offers complementary insights, rather than conflicting results. The OPP data confirmed that translation is acutely downregulated in TNF-α-treated cells, reflecting rapid suppression of mRNA translation. However, dynamic SILAC revealed that translation suppression is not uniform. While overall synthesis rates declined and cytosolic ribosomal proteins are degraded in atrophying conditions, mitochondrial proteins continued or even increased synthesis in late atrophy. This suggests that skeletal muscle cells selectively maintain the production of essential proteins to sustain metabolic function. Considering ATP’s regulatory role in protein synthesis (Buttgereit and Brand, 1995), short-term translation suppression may serve as an adaptive response saving ATP under catabolic conditions. Ribosomal dynamics further evolved over time: early atrophy showed cytoplasmic ribosome degradation and possible compensatory ribosome biogenesis, likely in response to ROS production (Sriram *et al*., 2011). In late atrophy, cytoplasmic ribosome synthesis stopped, potentially as an energy-saving strategy. Notably BafA1 treatment did not lead to an accumulation of known ribophagy receptors, suggesting alternative ribosome degradation mechanisms. Conversely, we observed increased mitochondrial ribosome synthesis in late atrophy, likely supporting OXPHOS and ATP production under catabolic stress, consistent with mitochondrial dysfunction reported in disuse atrophy (Cannavino *et al*., 2014).

To assess autophagic dynamics, we quantified LC3-II levels using a flow cytometry-based approach (Alsaleh *et al*., 2020). Autophagic turnover was increased at 24 hours of TNF-α treatment compared to homeostasis both by flow cytometry and in our SILAC datasets, corroborating findings of earlier studies using other techniques (Keller *et al*., 2011; Bernacchioni *et al*., 2021).

Under normal conditions, misfolded proteins are shuttled to the ER and degraded by the proteasome via ER-associated degradation (ERAD). Upon elevated stress, the accumulation of misfolded proteins can exceed ERAD capacity, leading to unfolded protein response (UPR) activation, which attempts to restore ER homeostasis by attenuating global protein synthesis (Harding, Zhang and Ron, 1999), promoting degradation of misfolded proteins via autophagy (Kouroku *et al*., 2007), and increasing chaperone expression (Yamamoto *et al*., 2007). Prolonged ER stress ultimately leads to a decline in protein folding capacity, likely exacerbating proteostasis disruptions and muscle damage (Bohnert *et al*., 2016). TNF-α has been shown to induce the UPR (Xue *et al*., 2005), and ATF4 activation in muscle atrophy models (Ebert *et al*., 2012) supporting the link between UPR and autophagy-mediated muscle atrophy. The chaperone Hspa5/BiP, crucial for ER protein folding, thereby maintaining ER homeostasis, accumulated upon BafA1 treatment in early atrophy compared to homeostasis, indicating its autophagic degradation early in atrophy. Whilst Hspa5/BiP is typically targeted for degradation via the UPS (Chang *et al*., 2016), we propose autophagic degradation under TNF-α-induced atrophy under severe or prolonged ER stress. Autophagic degradation of ER chaperones like Hspa5/BiP has been observed before upon proteasomal inhibition in *in vitro* studies by (Cha-Molstad *et al*., 2016). While early atrophy induced synthesis of receptors like Optn (aggrephagy, mitophagy) and Sec62 (ER-phagy), no ER-phagy receptors were newly synthesized at 72 hours. However, during late atrophy, we could observe significant accumulation of ER-phagy receptors Sec62, Calcoco1, and Tex264 – which are understudied ER-phagy receptors in the muscle context – as well as p62/Sqstm1 which acts as a promiscuous receptor in both canonical and non-canonical autophagy. Further supporting a progressive shift in proteostasis, protein folding pathways became increasingly downregulated in late atrophy. We could observe that TNF-α-induced atrophy shifts autophagic selectivity from maintaining homeostatic balance toward increased degradation of ER components upon atrophy stimulus.

Patients suffering from Duchenne muscular dystrophy (DMD) – a muscular disease caused by the loss of dystrophin (Hoffman, Brown and Kunkel, 1987) – eventually develop muscle atrophy due to a combination of interconnected factors. Chronic inflammation (Grounds and Torrisi, 2004) contributes to fibrosis, increased susceptibility to mechanical damage (Petrof *et al*., 1993) and impaired regeneration (Heslop, Morgan and Partridge, 2000), all of which progressively lead to muscle degeneration. DMD patients show an activation of the UPR and increased abundance of Hspa5/BiP (Moorwood and Barton, 2014; Pauly *et al*., 2017; Krishna *et al*., 2023) and a zebrafish DMD KO model with an Hspa5/BiP inhibitor improved muscle function (Ruparelia *et al*., 2024). Ruparelia *et al*. have also recently shown in zebrafish skeletal muscle that upon loss of Atrogin-1, a well-studied upregulated E3 ligase under atrophying conditions (Bodine *et al*., 2001; Bodine and Baehr, 2014) not only Hspa5/BiP accumulates but also leads to downregulation of autophagic flux in skeletal muscle impaired mitochondrial dynamics and loss of skeletal muscle fiber structure (Saneyasu *et al*., 2022). Upregulation of ER-stress/UPR-related factors and Hspa5/BiP was studied in a number of other muscle diseases like tibial muscular dystrophy caused by mutations in titin with altered autophagy levels (Screen *et al*., 2014). The progressive increase in ER degradation after TNF-α induced atrophy observed in our dynamic SILAC data, aligns with findings from Atg7 muscle knockout models where atrophy is exacerbated, accompanied by dilated sarcoplasmic reticulum (SR)/ER structures and observed upregulation of Hspa5/BiP, indicating activation of the UPR (Masiero *et al*., 2009).

Early atrophy involves upregulation of ubiquitin ligase activity, neddylation, and proteasomal function, indicating UPS activation. This was supported by the accumulation of myofibrillar proteins upon UPS inhibition across conditions. Late atrophic phase is marked by a pronounced degradation of myofibrillar proteins in our dynamic SILAC dataset as extensively previously characterized (Solomon and Goldberg, 1996), but also degradation of ECM and cytoskeletal components. This occurs alongside a decline in proteasome activity and neddylation, as indicated by GO terms associated with the significantly degraded proteins, suggesting a progressive impairment of the UPS over time. Despite increased autophagic turnover in atrophy, autophagy minimally contributes to myofibrillar degradation. No spatial localization of autophagy machinery to the M band was observed, even in the presence of p62/Sqstm1. Non-canonical functions for p62/Sqstm1 have been described before, linking it to activation of the Keap1-Nrf2 pathway for antioxidant response, (Komatsu *et al*., 2010; Sánchez-Martín *et al*., 2020), activation of mTORC1 in nutrient sensing (Duran *et al*., 2011) and activation of NF-κB during apopotosis and inflammation (Sanz *et al*., 2000).

The SR acts as a specialized form of the ER in skeletal muscle, regulating calcium storage and release for muscle contraction (Endo, Tanaka and Ogawa, 1970; Ford and Podolsky, 1970). Despite its distinct function, it retains core ER properties, including protein folding and processing. The fiber-type transition associated with inflammation-induced atrophy, from fast-twitch (glycolytic) to slow-twitch (oxidative) fibers (Mendell and Engel, 1971; Acharyya *et al*., 2004) may necessitate structural SR remodeling, given that glycolytic fibers have a more extensive SR network for rapid calcium cycling (Heilmann and Pette, 1979) while oxidative fibers rely more on mitochondrial ATP production. This aligns with our observation that mitochondrial ribosomal proteins are newly synthesized in atrophy, suggesting that muscle cells shift toward mitochondrial respiration phenotype as part of their adaptive response. Similar SR remodeling has been observed during exercise adaptation (Heilmann and Pette, 1979). Our results hint towards a highly regulated form of SR/ER remodelling via elevated autophagic activity beyond UPR during TNF-α-induced muscle atrophy.

Given our findings, further studies are warranted to explore whether inflammation similarly impairs ER-phagy in skeletal muscle, potentially contributing to the progression of age-related muscle wasting. While our dynamic SILAC study focuses specifically on TNF-α as an atrophic stimulus and utilizes the C2C12 in vitro model at defined time points, it offers valuable mechanistic insights into inflammation-induced muscle atrophy. Extending these findings to primary muscle cells and in vivo models will be key to fully capturing the complexity of muscle wasting. Nonetheless, this study highlights the potential of targeting autophagy and associated pathways, such as proteostasis regulation, mitochondrial remodeling, and ER-phagy, as novel therapeutic strategies in muscle atrophy.

## Material and Methods

### Cell culture

C2C12 mouse skeletal muscle myoblasts (ATCC # CRL-1772) were grown adherently and undifferentiated in DMEM (Sigma Aldrich # D5796) supplemented with 10% heat inactivated fetal bovine serum (FBS) (Gibco # 10500-064) and 1% Pen/Strep (P/S) (Sigma Aldrich # P4333). Myogenic differentiation was initiated upon reaching confluence by switching the cells to medium containing 2% FBS and kept in culture for seven to ten days, replenishing the medium every second day. Atrophy was induced on the seventh day with 10 ng/ml tumor necrosis factor α (TNF-α) (ThermoFisher # RMTNFAI) and cells cultivated for up to 72 h. For dynamic SILAC studies, cells were cultured in SILAC medium (PANBiotech # P04-02505) with 10% dialyzed FBS (Merck # F0392) and 200 mM GlutamaxTM (ThermoFisher 35050061). Cells were differentiated for seven to ten days in light SILAC medium (PANBiotech # P04-02505) supplemented with 200 mM GlutamaxTM (ThermoFiisher 35050061), 2% FBS, P/S (Gibco # 15140-122) and 28 mg/L light arginine 0 (R0 = Arginine 0 [^12^C_6_, ^14^N_4_], Sigma # A6969) and 48.7 mg/L light lysine (K0 = Lysine 0 [^12^C_6_, ^14^N_2_] Sigma # L8662). For pulsing with heavy medium, Arg0 and Lys0 in SILAC medium were replaced with 34 mg/l heavy arginine and 50.7 mg/l lysine (R10 = Arginine 10 [^13^C_6_, ^15^N_4_], Silantes 211604302 LOT (211CXN-LysI-503-02); K8 = Lysine 8 [^13^C_6_, ^15^N_2_], Silantes 201604102 LOT (201CXN-ArgI-603-01).

### Immunocytochemical (ICC) staining

Cells were cultured on ibidi polymer coverslips (Ibidi 81156), fixed in 4% paraformaldehyde (PFA) and permeabilized with DPBS (Gibco # 14040-117) containing 0.4 % Triton X-100. Proteins were stained with the following commercial antibodies: rat monoclonal anti-LAMP1-FITC (BioLegend # sc-121606 1:1000), rabbit polyclonal anti-LC3B (Novus Biologicals # NB100-2220 1:500), mouse monoclonal anti-MyHC (Sigma-Aldrich # M4276 1:1000), rabbit polyclonal anti-p62 (Invitrogen # PA5-20839 1:1000), rabbit monoclonal anti-Ub K63 (Sigma-Aldrich # 05-1308 1:1000), rabbit polyclonal anti-WIPI2 (Atlas Antibodies # HPA019852 1:400), goat anti-mouse IgG (H+L) secondary antibody AF647 (ThermoFisher # A-21235) and goat anti-rabbit IgG (H+L) secondary antibody AF488 (Invitrogen # 10729174). Cells were imaged on a Nikon Ti inverted microscope equipped with a Yokogawa CSU-W1 spinning disk confocal scanner, a Plan Apo λ 100x/1.45 NA oil-immersion objective and an Andor-DU-888 EMCCD camera. Fluorophores were excited using 488 nm and 647 nm laser lines. Images were acquired at 1024 x 1024 resolution. Z-stacks were acquired with step sizes ranging from 0.2-0.5 µm depending on the sample and region of interest. All image acquisition settings (exposure time, laser power, camera gain) were kept constant across conditions. Image processing was performed using Fiji (ImageJ 2.9.0), with only linear adjustments to brightness and contrast applied equally to all images.

### O-propargyl-puromycin (OPP) assay

For OPP labeling experiments, differentiated and atrophying C2C12 myotubes were incubated with 20 µM OPP (ThermoFisher, # C10459) for 30 min in growth medium at 37°C in the incubator and then fixed and permeabilized. Cycloheximide-treated samples served as negative control. The OPP-incorporated proteins were then fluorescently labeled via a copper-catalyzed click reaction and immediately analyzed via confocal microscopy. OPP puncta were manually counted across 20 images, each containing over 200 cells, with statistical significance assessed using ANOVA and Tukey’s test.

### Pharmacological inhibitors

To reduce the frequency of undifferentiated myoblasts in C2C12 cell populations which were subjected to differentiation, cells were treated with 50 µM Cytosin-1-β-D-arabinofuranosid (AraC, Sigma Aldrich # C1768) on day 4 of differentiation for 72 h. To inhibit proteolytic processes, either 50 µM Lactacystin (UPS inhibitor, Adoq Bioscience # A12768) or 50 nM Bafilomycin A1 (autophagy/lysosome inhibitor, Sigma Aldrich # B1793) was applied 3 h prior cell harvest. For translation inhibition, 50 µg/ml cycloheximide (Fluka BioChemika # 01811) was applied 90 min prior cell harvest or subsequent treatment.

### Automated protein processing and digestion for mass spectrometry

C2C12 cells were harvested, lysed in 50 mM Tris-HCl with 0.5% SDS and protease inhibitor (Roche, # 04693132001) and protein concentrations were determined using a BCA assay (Thermo Fisher Scientific #23250). Ten micrograms of each sample (100 µl) were transferred to a 96-well Armadillo plate and processed on an OT-2 robot (Opentrons). Proteins were reduced with 10 mM DTT at 37°C, 1000 rpm for 30 min, followed by alkylation with 15 mM iodoacetamide under the same conditions. The reaction was quenched with 30 mM DTT. Magnetic Sera-Mag SpeedBeads (1:20 bead-to-sample ratio) were added and incubated for 5 min at 1000 rpm. Proteins were bound with 100 µl acetonitrile, then washed sequentially with 80% ethanol and acetonitrile. Digestion was performed overnight at 37°C, 1150 rpm using LysC and trypsin (1:50 enzyme-to-protein ratio) in 100 mM ammonium bicarbonate. Peptides were collected into a new 96-well plate, dried via vacuum centrifugation and resuspended in Buffer A for LC-MS analysis.

### Mass spectrometry data acquisition

Each sample (1 µg) was analyzed using an Exploris 480 mass spectrometer coupled to a Vanquish Neo UHPLC (ThermoFisher) with a 106-min nanoflow (0.25 µl/min) gradient. Peptides were separated on a 20 cm, 1.9 µm in-house packed column. The gradient ramped from 2% to 30% of a buffer (0.1% formic acid, 90% acetonitrile) over 88 min, followed by 60% for 10 min, and 90% for 5 min. MS1 scans (350–1650Jm/z) were acquired at 120,000 resolution; MS2 scans used 40 isolation windows with a maximum injection time of 54 ms. The method was adapted from (Pekayvaz *et al*., 2022).

### Mass spectrometry data processing and analysis

Raw data were processed using DIA-NN (v1.8.1) with a project-specific spectral library generated from the *Mus musculus* UniProt FASTA database (UP000000589). SILAC-specific settings were applied (see Supplementary Methods for full parameters). Peptide-level quantification and SILAC ratio calculation were performed using the *silac_dia_tools_1.0*pipeline (https://github.com/rkerrid/silac_dia_tools_1.0). Light and heavy peptide intensities were extracted from DIA-NN outputs and used to compute direct (light/heavy), precursor-translated, and MS1-translated SILAC ratios. Precursor-level values were aggregated to the protein level using a log_2_-median approach. Protein intensities were normalized across runs using the directLFQ algorithm (https://github.com/MannLabs/directlfq). Data were log_2_-transformed prior to filtering. Proteins detected in fewer than two replicates or classified as contaminants were excluded. Missing or low-confidence values (e.g., intensities <0.001 or infinite) were set to NaN. Log_2_-transformed values were reverse-transformed where appropriate to maintain interpretability. Imputation was performed using the *silac_dia_statistics* pipeline (https://github.com/rkerrid/silac_dia_statistics), combining global distribution modeling with context-aware adjustments based on group-level sample minima. Imputed values were drawn from a normal distribution centered on the lower of the global shifted mean or the protein-specific minimum. Heavy-labeled intensities were further normalized via column-wise median-non-zero normalization.

### Statistical analysis

Differential abundance analysis was performed using the limma package in R (Ritchie *et al*., 2015) for all comparisons between datasets. Two-sample moderated t-tests were employed to assess differences in protein abundance between experimental conditions. To control the false discovery rate (FDR), p-values were adjusted using the Benjamini-Hochberg procedure. Proteins were classified as significantly degraded if the log_2_FC in the light channel was < 0 with an adj. p-value ≤ 0.05, significantly newly synthesized if log_2_FC in the heavy channel was > 0 with an adj. p-value ≤ 0.05 and significantly accumulated if the log_2_FC in the light channel was > 0 with an adj. p-value ≤ 0.05.

Figure 1B: Calculation of intensity clusters: Normalized light and heavy intensities were summed to calculate the total intensities of individual proteins at time points t0, t24, and t72 to create temporal profile clusters (Figure 1B). Cluster 2 was calculated as follows: constant intensity from t0 to t72, defined as exhibiting no more than a 10% variation relative to the maximum change intensity (Δ_max, with the 10% threshold calculated as 0.1 × Δ_max).

### Gene Ontology (GO) enrichment analysis

GO enrichment analysis (biological terms) was performed using Metascape to identify significantly overrepresented biological processes among differentially expressed proteins (Zhou *et al*., 2019).

### Quantification of LC3 using flow cytometry

C2C12 myotubes were differentiated for 7 days. On day 7, atrophy was induced by addition of 10 ng/ml TNF-α for 24 h or 72 h. Cells were harvested and washed three times with PBS. For flow cytometry analysis, cells were first stained with live/dead dye (Zombie NIR^TM^ Fixable Viability Kit, Biolegend # 423105). Afterwards, the Guava Autophagy LC3 antibody-based assay (Cytek # FCCH100171) was used to stain for LC3. Cells were permeabilized with PBS and 0.05% Saponin and cytosolic LC3 (LC3-I) was washed out before incubating with anti-LC3 antibody to only detect LC3-II. After the staining, cells were fixed with 4% PFA and data was acquired on a LSRFortessa^TM^ Flow cytometer (BD). FlowJo v10.10.0 was used for data analysis. For quantification of membrane-bound LC3-II, the geometric mean of the staining was quantified on live singlet cells.

## Author Credits

Conceptualization: Ursula K. Dueren, Matthias Selbach, Anna Katharina Simon

Data curation: Ursula K. Dueren, Alan An Jung Wei, A. Elisabeth Gressler, Oliver Popp

Formal analysis: Ursula K. Dueren, Alan An Jung Wei, A. Elisabeth Gressler, Oliver Popp

Investigation: Ursula K. Dueren, Alan An Jung Wei, A. Elisabeth Gressler

Methodology: Ursula K. Dueren, Alan An Jung Wei

Project administration: Ursula K. Dueren, Anna Katharina Simon

Validation: Ursula K. Dueren, A. Elisabeth Gressler

Visualization: Ursula K. Dueren, Anna Katharina Simon

Supervision: Matthias Selbach, Anna Katharina Simon, Thomas Sommer

Funding Acquisition: Thomas Sommer

Writing – original draft: Ursula K. Dueren, A. Elisabeth Gressler, Anna Katharina Simon

Writing – review and editing: Ursula K. Dueren, A. Elisabeth Gressler, Paolo Grumati, Matthias Selbach, Anna Katharina Simon, Thomas Sommer

## Conflict of Interest

The authors declare that they have no conflict of interest with the content of this article.

### Acknowledgements

This work was funded by the Max Delbrück Center for Molecular Medicine (MDC) in the Helmholtz Association and the MDC Graduate School. We acknowledge Simon Rapp and Anabel Y. Schweitzer for critical proofreading of the manuscript. We thank the light microscopy facility and the FACS facility of the MDC for their excellent technical support.

## Notes

### Competing Interest Statement

The authors have declared no competing interest.

