## Supplementary figures for "Autophagy selectively clears ER in inflammation-induced muscle atrophy"

### Figure S1

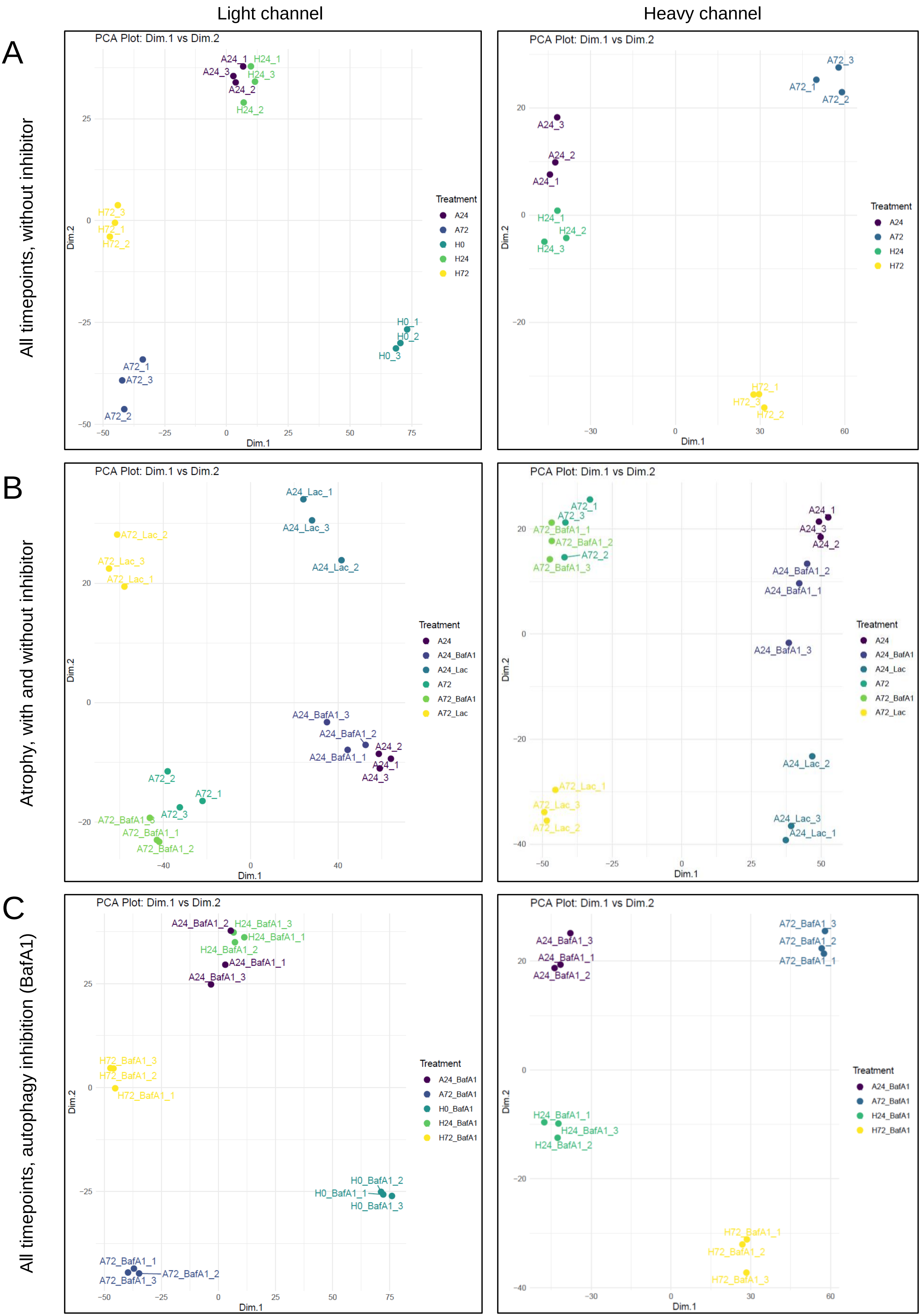

Light channel
Heavy channel
A
B
C
All timepoints, without inhibitor
Atrophy, with and without inhibitor
All timepoints, autophagy inhibition (BafA1)
PCA Plot: Dim.1 vs Dim.2
Dim.1
Dim.2
Treatment
A24
A72
H0
H24
H72
A24_1
A24_2
A24_3
H24_1
H24_2
H24_3
H72_1
H72_2
H72_3
A72_1
A72_2
A72_3
H0_1
H0_2
H0_3
A24_BafA1
A24_Lac
A72_BafA1
A72_Lac
A24_Lac_1
A24_Lac_2
A24_Lac_3
A72_Lac_1
A72_Lac_2
A72_Lac_3
A24_BafA1_1
A24_BafA1_2
A24_BafA1_3
A72_BafA1_1
A72_BafA1_2
A72_BafA1_3
H0_BafA1
H24_BafA1
H72_BafA1
H0_BafA1_1
H0_BafA1_2
H0_BafA1_3
H24_BafA1_1
H24_BafA1_2
H24_BafA1_3
H72_BafA1_1
H72_BafA1_2
H72_BafA1_3

### Figure S2

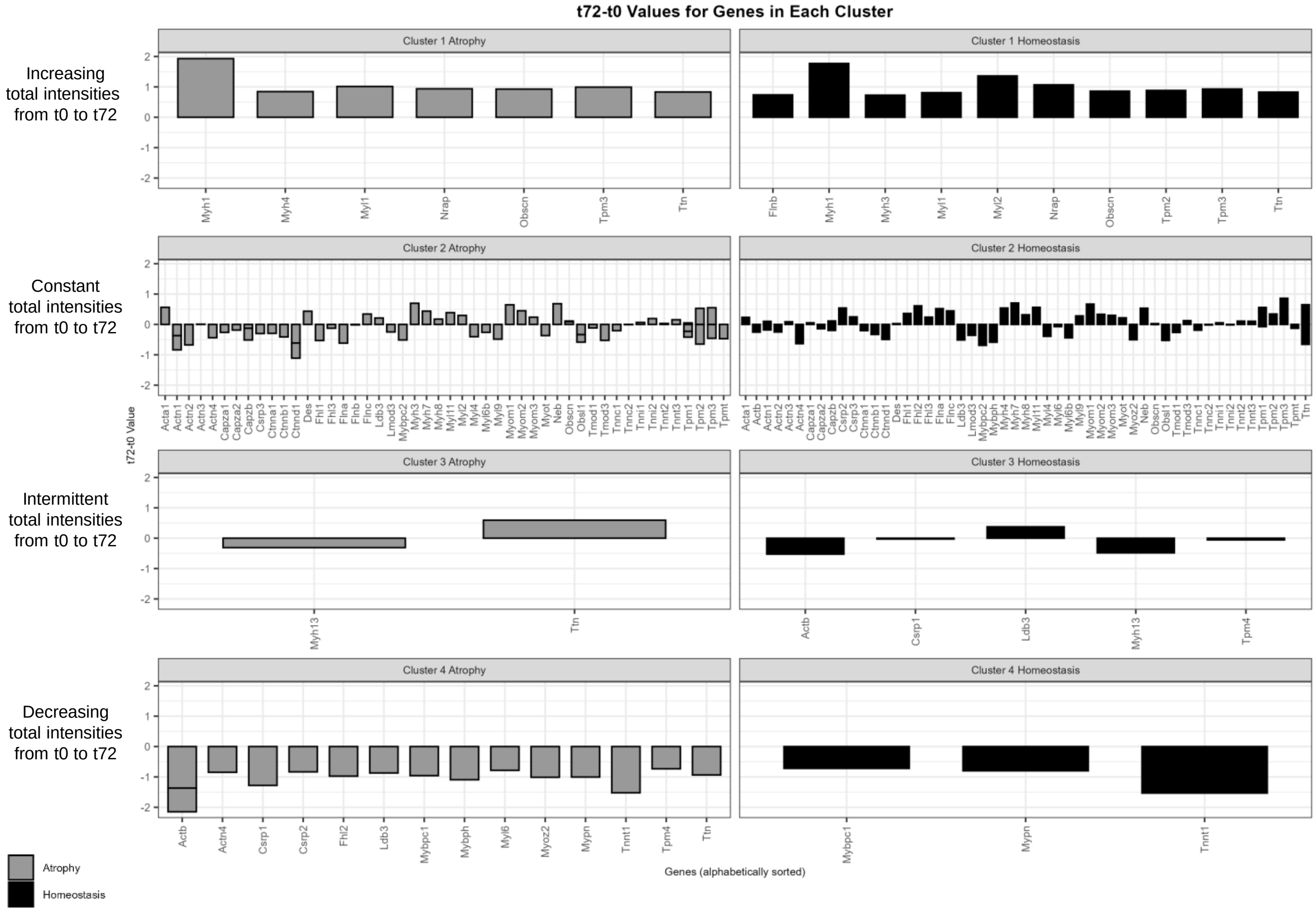
t72-t0 Values for Genes in Each Cluster
Increasing total intensities from t0 to t72
Constant total intensities from t0 to t72
Intermittent total intensities from t0 to t72
Decreasing total intensities from t0 to t72
Cluster 1 Atrophy
Cluster 1 Homeostasis
Myh1 Myh4 Myl1 Nrap Obscn Tpm3 Ttn
Flnb Myh1 Myh3 Myl1 Myl2 Nrap Obscn Tpm2 Tpm3 Ttn
Cluster 2 Atrophy
Cluster 2 Homeostasis
Cluster 3 Atrophy
Cluster 3 Homeostasis
Myh13 Ttn
Actb Csrp1 Ldb3 Myh13 Tpm4
Cluster 4 Atrophy
Cluster 4 Homeostasis
Actb Actn4 Csrp1 Csrp2 Fhl2 Ldb3 Mybpc1 Mybph Myl6 Myoz2 Mypn Tnnt1 Tpm4 Ttn
Mybpc1 Mypn Tnnt1
t72-t0 Value
Genes (alphabetically sorted)
Atrophy
Homeostasis

Figure S3

A

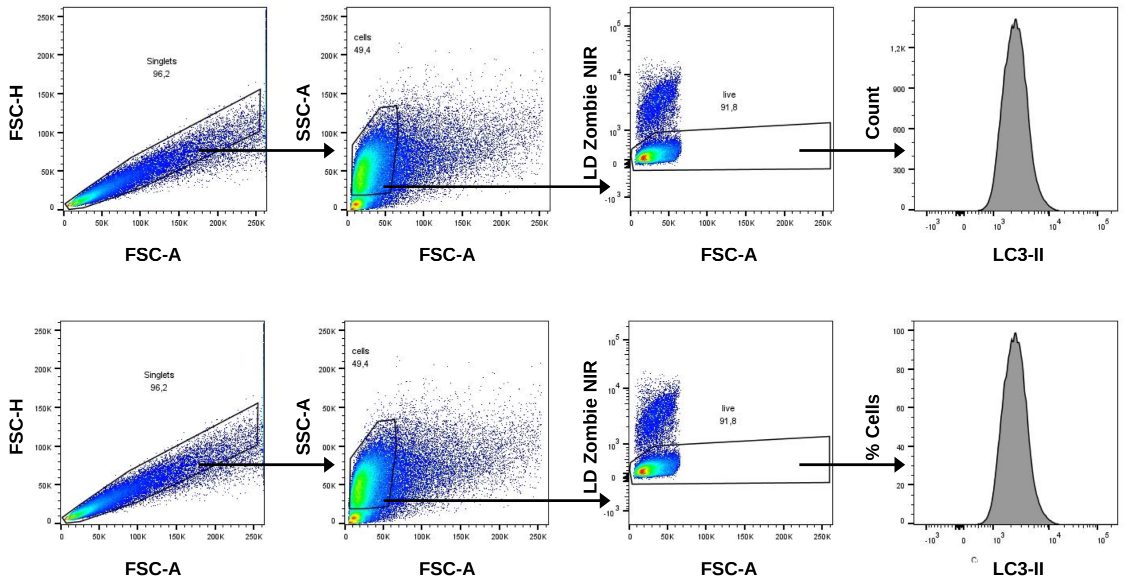
FSC-H
Singlets
96,2
FSC-A
SSC-A
cells
49,4
FSC-A
LD Zombie NIR
live
91,8
FSC-A
Count
LC3-II
FSC-H
Singlets
96,2
FSC-A
SSC-A
cells
49,4
FSC-A
LD Zombie NIR
live
91,8
FSC-A
% Cells
LC3-II

### Figure S4

**Homeostasis (H24)**

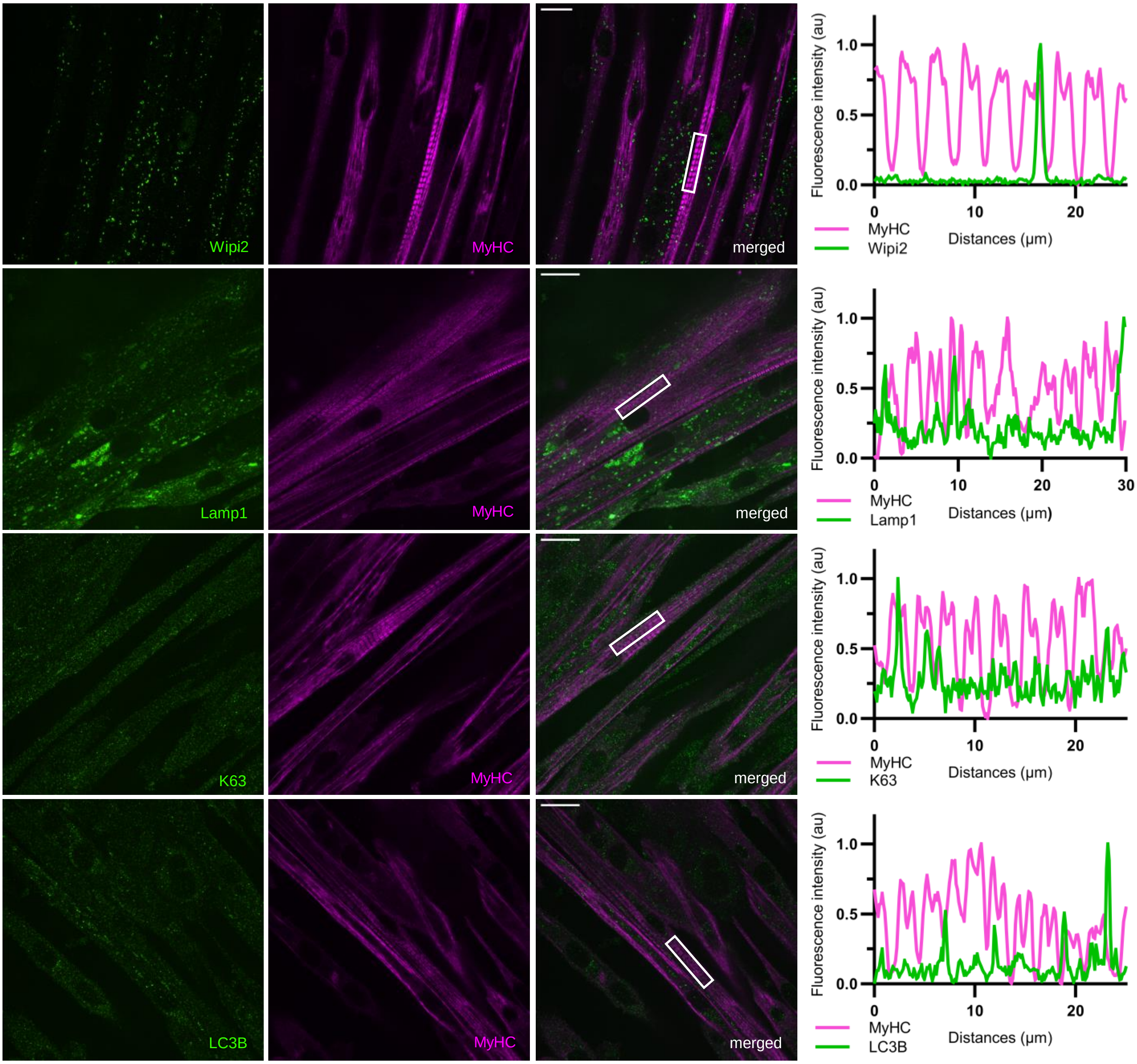

### Figure S5

**Atrophy 24 h (A24)**

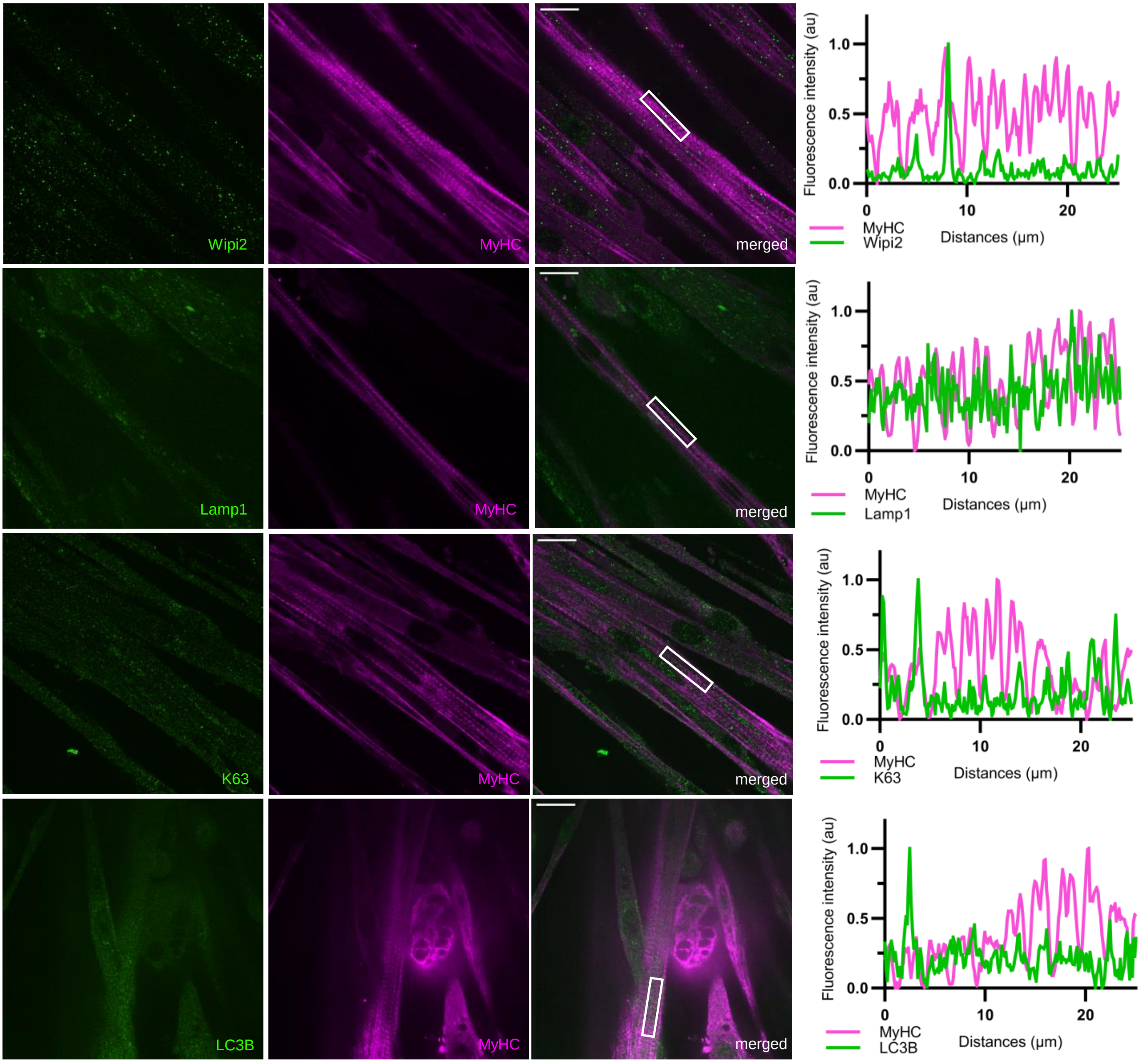

### Figure S6

A

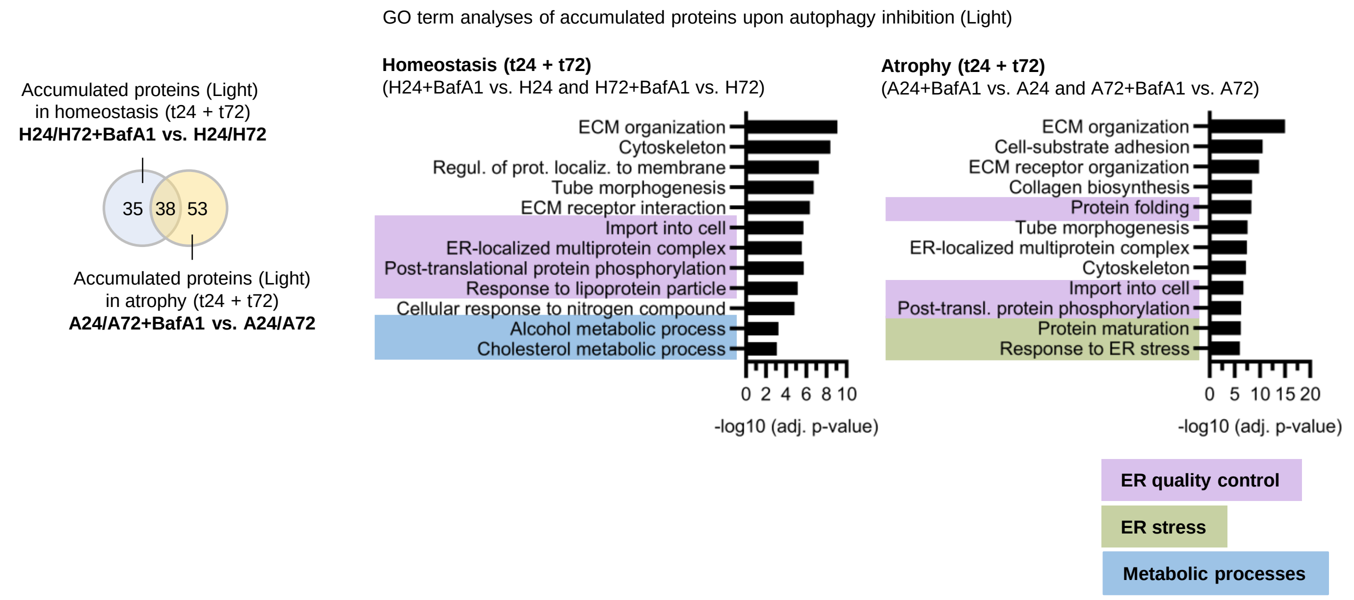
Accumulated proteins (Light)
in homeostasis (t24 + t72)
H24/H72+BafA1 vs. H24/H72
35
38
53
Accumulated proteins (Light)
in atrophy (t24 + t72)
A24/A72+BafA1 vs. A24/A72
GO term analyses of accumulated proteins upon autophagy inhibition (Light)
Homeostasis (t24 + t72)
(H24+BafA1 vs. H24 and H72+BafA1 vs. H72)
ECM organization
Cytoskeleton
Regul. of prot. localiz. to membrane
Tube morphogenesis
ECM receptor interaction
Import into cell
ER-localized multiprotein complex
Post-translational protein phosphorylation
Response to lipoprotein particle
Cellular response to nitrogen compound
Alcohol metabolic process
Cholesterol metabolic process
0 2 4 6 8 10
-log10 (adj. p-value)
Atrophy (t24 + t72)
(A24+BafA1 vs. A24 and A72+BafA1 vs. A72)
ECM organization
Cell-substrate adhesion
ECM receptor organization
Collagen biosynthesis
Protein folding
Tube morphogenesis
ER-localized multiprotein complex
Cytoskeleton
Import into cell
Post-transl. protein phosphorylation
Protein maturation
Response to ER stress
0 5 10 15 20
-log10 (adj. p-value)
ER quality control
ER stress
Metabolic processes

B

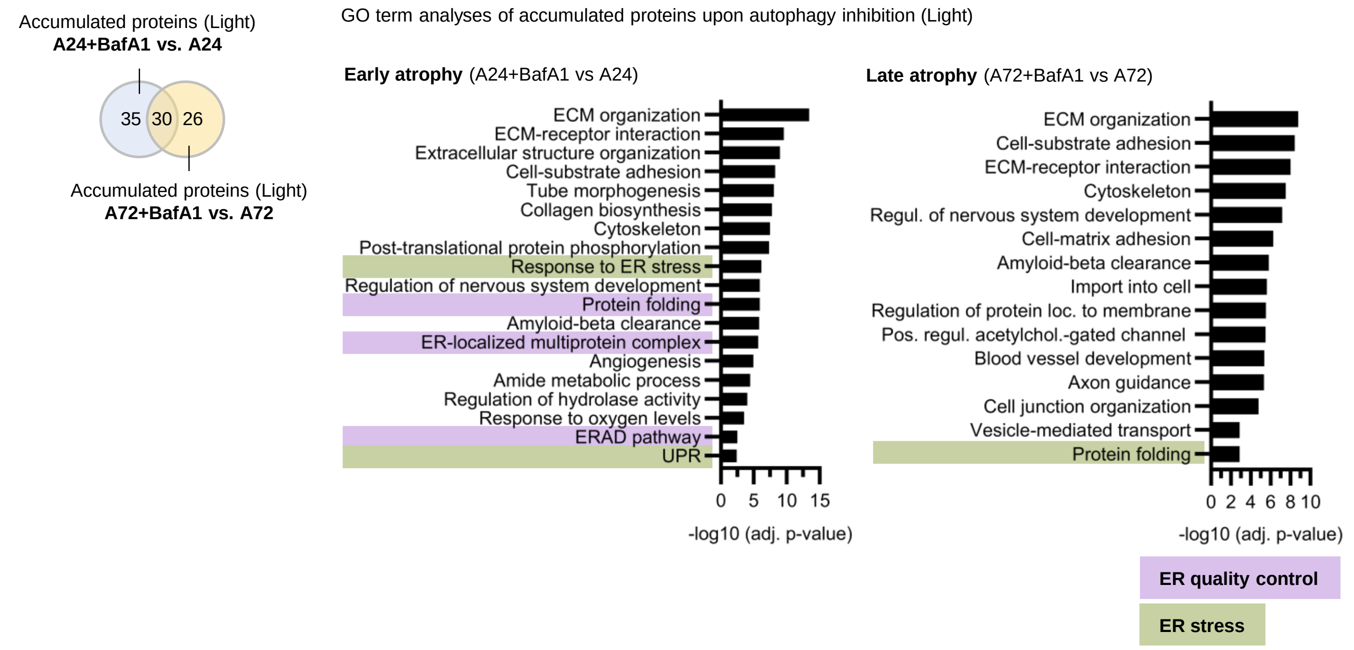
Accumulated proteins (Light)
A24+BafA1 vs. A24
35
30
26
Accumulated proteins (Light)
A72+BafA1 vs. A72
GO term analyses of accumulated proteins upon autophagy inhibition (Light)
Early atrophy (A24+BafA1 vs A24)
ECM organization
ECM-receptor interaction
Extracellular structure organization
Cell-substrate adhesion
Tube morphogenesis
Collagen biosynthesis
Cytoskeleton
Post-translational protein phosphorylation
Response to ER stress
Regulation of nervous system development
Protein folding
Amyloid-beta clearance
ER-localized multiprotein complex
Angiogenesis
Amide metabolic process
Regulation of hydrolase activity
Response to oxygen levels
ERAD pathway
UPR
0 5 10 15
-log10 (adj. p-value)
Late atrophy (A72+BafA1 vs A72)
ECM organization
Cell-substrate adhesion
ECM-receptor interaction
Cytoskeleton
Regul. of nervous system development
Cell-matrix adhesion
Amyloid-beta clearance
Import into cell
Regulation of protein loc. to membrane
Pos. regul. acetylchol.-gated channel
Blood vessel development
Axon guidance
Cell junction organization
Vesicle-mediated transport
Protein folding
0 2 4 6 8 10
-log10 (adj. p-value)
ER quality control
ER stress

### Figure S7

## A

Accumulated proteins (Light)
in homeostasis (t24 + t72)
**H24/H72+BafA1 vs. H24/H72**

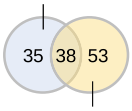


Accumulated proteins (Light)
in atrophy (t24 + t72)
**A24/A72+BafA1 vs. A24/A72**

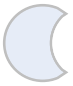

| | |
|---|---|
| Acadl | Mtch2 |
| Acsl3 | Ndufa10 |
| Aldh3a2 | Nol6 |
| Calu | Npc1 |
| Ccar1 | Pcbd2 |
| Colgalt1 | Plscr3 |
| Crybg2 | Porcn |
| Csk | Rab33b |
| Dnajc5 | Sgca |
| Dtymk | Slain2 |
| Ephb2 | Slc12a2 |
| Fahd2a | Slc25a4 |
| Fam171a2 | Tmem263 |
| Fat1 | Tmem63a |
| Hk1 | Uggt1 |
| Hyou1 | Vamp2 |
| Jup | |
| Kct2 | |
| Minpp1 | |

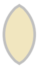

| | |
|---|---|
| App | Ldlr |
| Bgn | Lrp1 |
| Calcoco1 | Lrrn1 |
| Cdh13 | Map1lc3a |
| Cdh15 | Mesd |
| Chrnb1 | Mfge8 |
| Col1a2 | Msln |
| Cpd | Musk |
| Dag1 | Nid2 |
| Ephb3 | Plxnb2 |
| Fam234a | Ranbp2 |
| Fn1 | Serinc1 |
| Gpc1 | Serpinh1 |
| Hsp90b1 | Sparc |
| Hspg2 | Sqstm1 |
| Igf2r | Stx12 |
| Itga7 | Tfrc |
| Itm2b | Thbs1 |
| Jam2 | Timp1 |

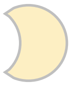

| | | |
|---|---|---|
| Acot2 | H4c1 | Pdia4 |
| Antxr2 | Hspa5 | Plod1 |
| Atp13a3 | Ifitm3 | Plod3 |
| Atp6ap1 | Ilf2 | Poglut2 |
| Atp6v1d | Itga6 | Rab11b |
| B2m | Lamb2 | Rdh14 |
| Bcam | Lnpep | Sar1a |
| Cemip2 | Lrp4 | Scfd2 |
| Col12a1 | Magoh | Sgpl1 |
| Dnajc3 | Ncam1 | Slc25a40 |
| Ece1 | Nid1 | Slc29a1 |
| Epha2 | Nomo1 | Slc3a2 |
| Erbb3 | Npnt | Slc7a5 |
| Exosc4 | Nt5dc2 | Snrpb2 |
| F11r | Nup85 | Tor1a |
| Flot1 | P4hb | |
| Fzd7 | Pcolce | |
| Gabarapl2 | Pcyox1 | |
| Gpr107 | Pdia3 | |

## B

Accumulated proteins (Light)
**A24+BafA1 vs. A24**

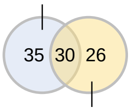


Accumulated proteins (Light)
**A72+BafA1 vs. A72**

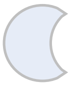

| | |
|---|---|
| Acot2 | Nup85 |
| Antxr2 | P4hb |
| Atp6ap1 | Pcyox1 |
| B2m | Pdia3 |
| Cemip2 | Plod1 |
| Col12a1 | Plod3 |
| Dnajc3 | Poglut2 |
| Epha2 | Sar1a |
| Erbb3 | Scfd2 |
| Exosc4 | Serpinh1 |
| Fam234a | Sgpl1 |
| Flot1 | Slc25a40 |
| Fzd7 | Slc29a1 |
| Hspa5 | Slc3a2 |
| Itga7 | Snrpb2 |
| Mfge8 | Timp1 |
| Npnt | Tor1a |
| Nt5dc2 | |

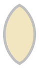

| | |
|---|---|
| App | Igf2r |
| Bgn | Itga6 |
| Calcoco1 | Itm2b |
| Cdh15 | Jam2 |
| Chrnb1 | Ldlr |
| Col1a2 | Lrp1 |
| Cpd | Lrp4 |
| Dag1 | Lrrn1 |
| Ece1 | Msln |
| Ephb3 | Nid2 |
| Fn1 | Plxnb2 |
| Gpc1 | Serinc1 |
| Hsp90b1 | Sparc |
| Hspg2 | Sqstm1 |
| Ifitm3 | Thbs1 |

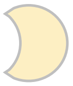

| | |
|---|---|
| Atp13a3 | Ncam1 |
| Atp6v1d | Nid1 |
| Bcam | Nomo1 |
| Cdh13 | Pcolce |
| F11r | Pdia4 |
| Gabarapl2 | Rab11b |
| Gpr107 | Ranbp2 |
| H4c1 | Rdh14 |
| Ilf2 | Slc7a5 |
| Lamb2 | Stx12 |
| Lnpep | Tfrc |
| Magoh | |
| Map1lc3a | |
| Mesd | |
| Musk | |

### Figure S8

A

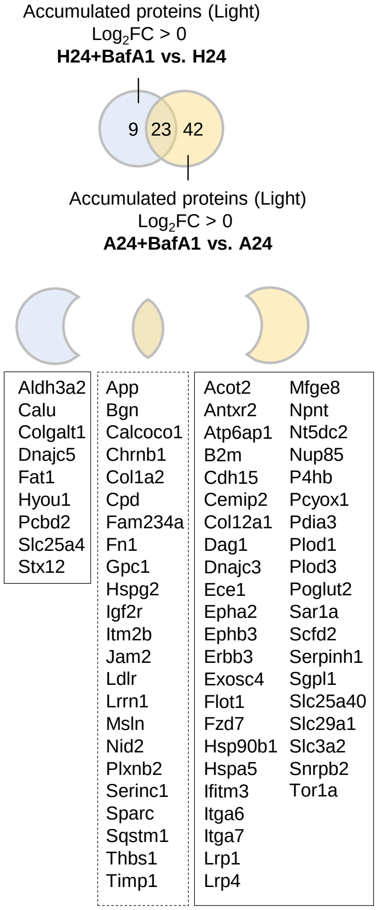


B

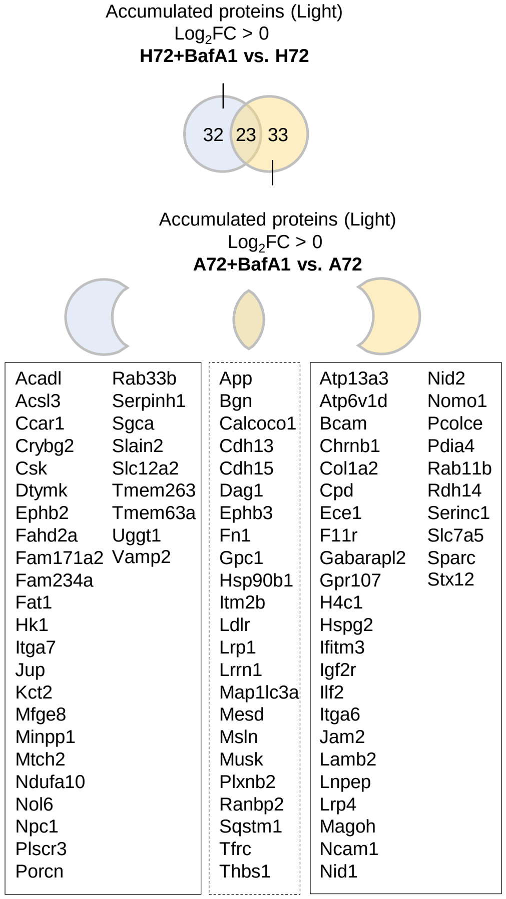


C

GO term analyses of accumulated proteins upon autophagy inhibition in homeostasis (Light)

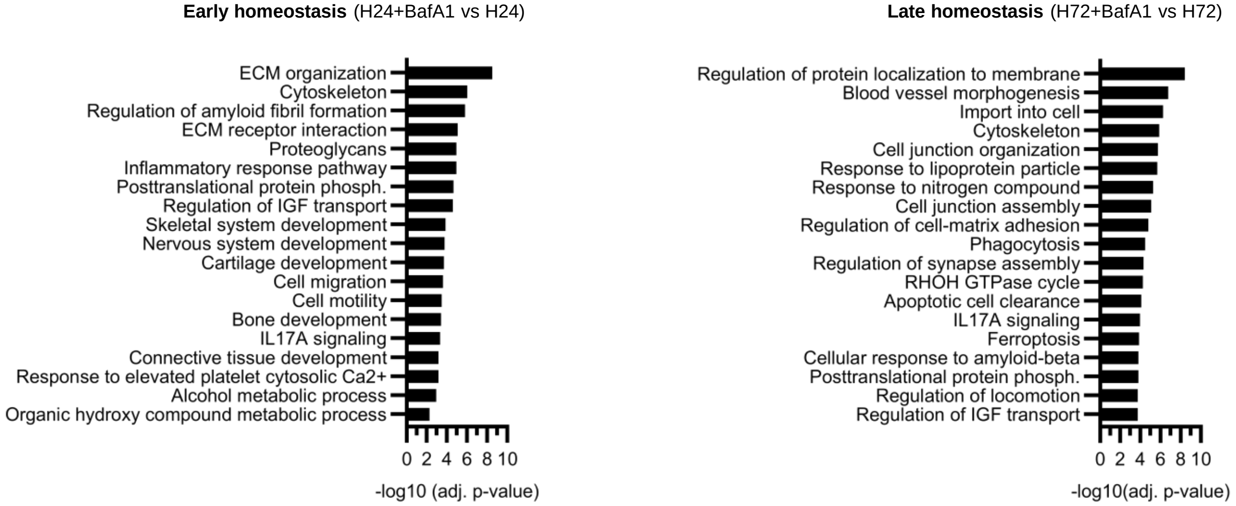

### Figure S9

A

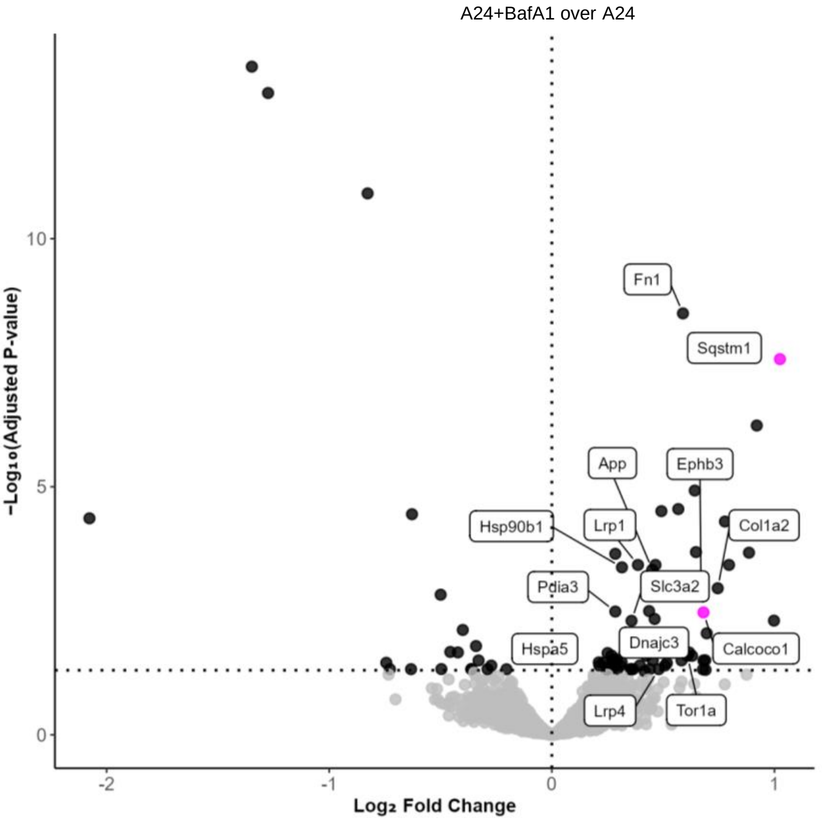

A24+BafA1 over A24
−Log₁₀(Adjusted P-value)
Log₂ Fold Change
Fn1
Sqstm1
App
Ephb3
Hsp90b1
Lrp1
Col1a2
Pdia3
Slc3a2
Hspa5
Dnajc3
Calcoco1
Lrp4
Tor1a


B

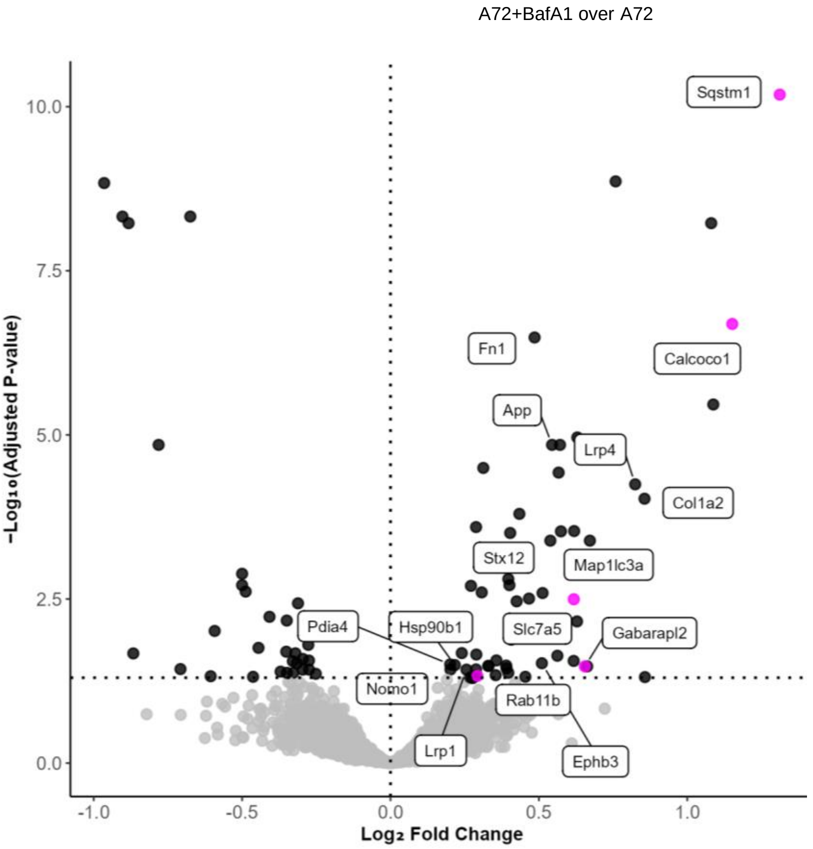

A72+BafA1 over A72
−Log₁₀(Adjusted P-value)
Log₂ Fold Change
Sqstm1
Fn1
Calcoco1
App
Lrp4
Col1a2
Stx12
Map1lc3a
Pdia4
Hsp90b1
Slc7a5
Gabarapl2
Nomo1
Rab11b
Lrp1
Ephb3
