## Supplementary text for "Autophagy selectively clears ER in inflammation-induced muscle atrophy"

### Supplementing Information Material and Methods

**DIA-NN parameters**

DIA-NN (version 1.8.1) was run with the following parameters to support dynamic SILAC quantification:

- --fixed-mod SILAC,0.0,KR,label
- --lib-fixed-mod SILAC
- --channels SILAC,L,KR,0:0;SILAC,H,KR,8.014199:10.008269
- --peak-translation
- --original-mods
- --no-norm
- --no-maxlfq
- --peak-center

Additional settings:

- Missed cleavages: 2
- Precursor charge range: 2–4
- Precursor m/z range: 300–1800
- Fragment ion m/z range: 200–1800

### Supplemental figure legends

**Figure S1:** PCA of dynamic SILAC dataset of light and heavy channels comparing **(A)** all time points without proteolytic inhibitors, **(B)** atrophying time points with and without proteolytic inhibitors and **(C)** all time points upon autophagy inhibition. A = atrophy, H = homeostasis, 24 = 24 hours, 72 = 72 hours, 0 = 0 hours with 1-3 depicting three replicates. BafA1 = BafilomycinA1, Lac = Lactacystin.

**Figure S2:** Myofibrillar proteins were assigned to clusters based on their temporal profile of total intensities (Heavy + Light) between t0, t24 and t72. Clusters were identified for atrophying (left) and homeostatic (right) conditions. Clusters are defined as: 1: increasing, 2: constant, 3: intermittent, 4: decreasing. Y-axis depicts t72-t0 value. Genes are alphabetically ordered within their clusters.

**Figure S3:** **(A)** Gating strategy for quantification of LC3-II intensity using flow cytometry. Autophagic turnover was quantified using a flow cytometry-based approach (Alsaleh *et al.*, 2020). For analysis, doublets were excluded first, followed by gating on cells, thus excluding debris and enlarged cells. Next, dead cells were removed by gating on live cells. LC3-II signal was thus only quantified on live, singlet cells. **(B)** Dynamic SILAC dataset: Subset of accumulated autophagic machinery proteins (Light) of ANOVA comparisons of Bafilomycin (BafA1)-treated atrophying (A24, A72) vs. homeostatic (H24, H72) cells at time point t24 and t72 with log_2_FC > 0 and adj. p-value set at < 0.05.

**Figure S4:** Immunocytochemical stainings for Wipi2, Lamp1, K63-Ub, and LC3B show no colocalization with MyHC (Myosin heavy chain) in myotubes with corresponding line plots along the sarcomere. Scale bar: 20 µm.

**Figure S5:** Immunocytochemical stainings for Wipi2, Lamp1, K63-Ub, and LC3B show no colocalization with MyHC (Myosin heavy chain) in atrophying myotubes (A24, 24 h TNF-α treatment) with corresponding line plots along the sarcomere. Scale bar: 20 µm.

**Figure S6: (A)** ANOVA comparisons of Bafilomycin (BafA1)- vs. non-BafA1-treated samples in homeostasis (both H24 and H72 together) and atrophy (both A24 and A72 together) with log_2_FC > 0 and adj. p-value set at < 0.05. Numbers in Venn diagrams depict amount of significantly regulated proteins in respective dataset comparison. GO term analyses (biological processes) in homeostasis (left) and atrophy (right). **(B)** ANOVA comparisons of Bafilomycin (BafA1)- vs. non-BafA1-treated samples in early atrophy (A24+BafA1 vs. A24) and late atrophy (A72+Baf A1 vs. A72) with log_2_FC > 0 and adj. p-value set at < 0.05. Numbers in Venn diagrams depict amount of significantly regulated proteins in respective dataset comparison. GO term analyses (biological processes) in early atrophy (left) and late atrophy (right).

**Figure S7:** **(A)** ANOVA comparisons of Bafilomycin (BafA1)- vs. non-BafA1-treated samples in homeostasis (both H24 and H72 together) and atrophy (both A24 and A72 together) with log_2_FC > 0 and adj. p-value set at < 0.05. Numbers in Venn diagrams depict amount of significantly regulated proteins in respective dataset comparison. Lists depict accumulated proteins in only homeostasis (left), only atrophy (right) and proteins accumulating in both conditions (overlap, middle). **(B)** ANOVA comparisons of Bafilomycin (BafA1)- vs. non-BafA1-treated samples in early atrophy (A24+BafA1 vs. A24) and late atrophy (A72+Baf A1 vs. A72) with log_2_FC > 0 and adj. p-value set at < 0.05. Numbers in Venn diagrams depict amount of significantly regulated proteins in respective dataset comparison. Lists depict accumulated proteins in only early atrophy (left), only late atrophy (right) and proteins accumulating in both conditions (overlap, middle).

**Figure S8:** ANOVA comparisons of Bafilomycin (BafA1)- vs. non-BafA1-treated samples in both homeostasis and atrophy at **(A)** 24 hours and **(B)** 72 hours with log_2_FC > 0 and adj. p-value set at < 0.05. Numbers in Venn diagrams depict amount of significantly regulated proteins in respective dataset comparison. Lists depict accumulated proteins in only homeostasis (left), only atrophy (right) and proteins accumulating in both conditions (overlap, middle). **(C)** GO term analyses (biological processes) of ANOVA comparisons of Bafilomycin (BafA1)- vs. non-BafA1-treated samples in homeostasis at 24 hours (left) and 72 hours (right).

**Figure S9:** Volcano plots depicting significantly accumulating proteins of ANOVA comparisons of BafA1- vs. non-BafA1-treated samples in early (A24) and late (A72) atrophy with magenta-coloured dots depicting autophagy markers. Log_2_FC >0 and adj. p-value set at < 0.05. BafA1 = BafilomycinA1.
